# Ancient and modern genomes reveal microsatellites maintain a dynamic equilibrium through deep time

**DOI:** 10.1101/2020.03.09.972364

**Authors:** Bennet J. McComish, Michael A. Charleston, Matthew Parks, Carlo Baroni, Maria Cristina Salvatore, Ruiqiang Li, Guojie Zhang, Craig D. Millar, Barbara R. Holland, David M. Lambert

## Abstract

Microsatellites are widely used in population genetics, but their evolutionary dynamics remain poorly understood. It is unclear whether microsatellite loci drift in length over time. We identify more than 27 million microsatellites using a novel and unique dataset of modern and ancient Adélie penguin genomes along with data from 63 published chordate genomes. We investigate microsatellite evolutionary dynamics over two time scales: one based on the Adélie penguin samples dating to approximately 46.5 kya, the other dating to the diversification of chordates more than 500 Mya. We show that the process of microsatellite allele length evolution is at dynamic equilibrium; while there is length polymorphism among individuals, the length distribution for a given locus remains stable. Many microsatellites persist over very long time scales, particularly in exons and regulatory sequence. These often retain length variability, suggesting that they may play a role in the maintenance of evolutionary plasticity.

## Introduction

Microsatellites or short tandem repeats (STRs), consisting of tandem repeats of two to six base pair motifs, are prevalent in both prokaryotic and eukaryotic genomes. Some microsatellites have been shown to be functionally important^1-3^, but most are assumed to evolve neutrally, and for this reason, along with their abundance and high variability, they have been used extensively in population genetics studies^4^. However, their evolutionary dynamics remain poorly understood, and it is unclear whether microsatellite loci are in dynamic equilibrium with respect to the length of alleles, or whether alleles experience directional drift in length. This is important because the mutation processes that underlie these important genetic markers are central to the evolutionary models that employ microsatellites.

In this study, when describing microsatellites, we consider both the number of base pairs in the underlying motif *period* and, for each allele, how many times the motif appears (the *repeat number*). We refer to microsatellites that contain only exact copies of the motif as *pure*. The total length of a microsatellite allele (in nucleotides) is the product of the period and repeat number. Repeat number is thought to change through a process of replication slippage^5,6^, by which strands may transiently dissociate during DNA replication and then mispair with a different copy of the repeat, resulting in the insertion or deletion of one or more repeat units. Microsatellites are highly plastic in evolutionary terms, with mutation rates due to replication slippage generally several orders of magnitude higher than for point mutation^7^.

Much still remains to be learned about the mutational processes involved in microsatellite evolution. The overall process can be thought of as a birth–death process (increase or decrease in length of microsatellite by the birth or death of individual repeat units) embedded within a second birth–death process (microsatellite loci appear and disappear) all happening along a branching process (the population history). Slippage during DNA replication is thought to be the main cause of changes in length, with mismatch repair reducing the mutation rate^8^, but recombination may also play a role, and point mutations must be taken into account. The processes by which new microsatellites appear, and by which they eventually degenerate and disappear, are particularly poorly understood^9^.

Existing models are highly simplified and only take into account changes in length (and occasionally purity) in existing microsatellites, ignoring the processes of ‘birth’ and ‘death’ by which microsatellite loci appear and disappear (and perhaps reappear)^10^. Some of these models have been designed so that they have a stationary distribution (for example those of Kruglyak *et al.*^11^, Calabrese *et al.*^12^ and Amos *et al.*^13^), but it is not clear whether this is biologically realistic. It may be that an individual microsatellite locus is never at equilibrium, tending instead to increase in length throughout its life, but that the birth-death process causes the genome-wide distribution of allele lengths at all microsatellite loci to be at equilibrium.

A key open question is thus whether the alleles at a microsatellite locus increase or decrease in average length over time, or whether each locus is maintained at an equilibrium length. While some pedigree studies have shown a bias in favor of gain of repeats^14^, suggesting that microsatellites should rapidly increase in size ^15^, others have found that slippage has a length-dependent bias^16-18^,supporting earlier suggestions that constraints exist on repeat number at microsatellite loci^19^. On the basis of the former observation, it has been suggested that microsatellites increase in length until the accumulation of point mutations hinders slippage and ultimately leads to the degeneration of the microsatellite locus^10,11^. Alternatively, Amos *et al.*^13^ recently proposed a model consistent with the latter observations, in which inter-allelic interactions in heterozygous individuals may drive the process whereby longer-than-average alleles tend to get shorter and shorter-than-average alleles tend to get longer (which they call the centrally directed mutation model).

Here we make use of exceptionally well-preserved ancient DNA from a unique set of Adélie penguin samples reported here for the first time, and genotype 177,974 microsatellites in both modern and ancient genomes, including some dating to approximately 46.5 kya. In addition, we are able to time the evolutionary origin of many of these loci by aligning them with more than 27 million microsatellites from a large set of published chordate genomes and mapping them onto a recent phylogeny^20^. Our data include microsatellites that date to the diversification of chordates more than 500 Mya. We show that allele lengths at microsatellite loci are in dynamic equilibrium, and these have remained stable over hundreds of millions of years and through many speciation events. While there is length polymorphism among individuals, the overall length distribution for a given locus does not change appreciably over time. We show that microsatellites can persist over very long time scales, particularly those in exons and regulatory sequence, while retaining length variability. This suggests that microsatellites may play a role in the maintenance of evolutionary plasticity.

## Results

### Microsatellite dynamics in Adélie penguin samples

Genomes obtained from ancient biological remains allow us to observe changes in sequence variation that cannot be observed using only contemporary sequences. Here we have used whole-genome sequence data of ancient Adélie penguin remains from 23 individuals dated at up to 46,587 years old, as well as from 26 modern individuals, to identify 177,974 microsatellite loci in an Adélie penguin reference genome. Most loci are close to the minimum length detectable for each period (especially in the case of pure loci), with very small numbers of longer loci up to thousands of base pairs in length. The length distributions of these microsatellite loci are shown in Supplementary Fig. 1. We determined the genotype of these loci in each of the ancient and modern Adélie samples, and allele length distributions for each sample are shown in Supplementary Fig. 2. These genotype data enable us to obtain length distributions for microsatellites at different time points, and hence to test whether there is any evidence for directional drift in microsatellite length.

To test whether microsatellite allele length is stationary, or whether the average allele lengths of individual loci increase over time, we used BayesFactor^21^ to compare generalized linear mixed models in which allele length is treated as dependent on different combinations of possible explanatory variables. The explanatory variables considered were: the motif of the allele, the surrounding sequence type (exon, intron, regulatory, or intergenic), and sample age. We also tested for an interaction between surrounding sequence type and sample age. In addition to these fixed effects, which are assumed to be the same for all genomes, we also treated the sample, i.e., the particular Adélie genome, as a random effect; this is equivalent to allowing a different intercept in the regression model for each genome. Impurity affects the length at which microsatellites can be detected, so models were fit separately for pure and impure microsatellites. Similarly, models were fit separately for loci of different periods because different alignment score thresholds were used to detect them, so that their allele lengths cannot be compared directly. Bayes factors for all models tested are given in Supplementary Table 1, and posterior estimates of effect sizes in Supplementary Table 2. For both pure and impure microsatellites of each period, the best-supported model is that in which allele length is dependent on the motif and surrounding sequence type. Our data provide positive evidence for this model, being at least seven times more likely to be observed under this model than under a model in which length depends on sample age. Since length does not depend on sample age in this model, we infer that the process of expansion and contraction of microsatellite alleles is effectively stationary, or nearly stationary, over a time-scale of tens of thousands of years.

### Microsatellite locus age inference

To investigate microsatellite dynamics over a much longer timescale and across a broad range of species, we used whole-genome alignments of 48 avian species from Zhang *et al.*^22^ along with the genomes of fifteen non-avian vertebrate species that span the chordate tree. We identified a total of over 27 million microsatellites in the 63 genomes, and a breakdown of the numbers of loci of each period detected in each genome is given in Supplementary Table 3. For each of these species, we used a whole-genome alignment to chicken to generate a standard set of coordinates for all microsatellites present in the alignment. The number of microsatellite loci in any species that can be aligned to the chicken genome, and the overall number of bases aligned to the chicken genome, are negatively correlated with the time since the most recent common ancestor of that species and chicken (see Supplementary Fig. 3). We were able to map approximately 5.4 million microsatellites across the 63 species to almost 2.9 million loci in the chicken genome. Of these, almost 2.2 million microsatellite loci were found in only a single species and 680,804 loci had microsatellites conserved across two or more species. Exact numbers of microsatellites detected and aligned are given in Supplementary Table 4.

We used the dated avian whole-genome phylogeny published by Jarvis *et al.*^20^, to which we added 15 non-avian species with estimated divergence times taken from the Timetree of Life (www.timetree.org)^23^. To infer gains and losses of microsatellite loci in different lineages, we carried out ancestral state reconstruction on a subtree whose topology is relatively uncontroversial, agreeing with the trees published by Jarvis *et al.*^20^ and Prum *et al.*^24^, and on which we expect incomplete lineage sorting events to be rare^25^. This allows us to infer the edge on which any locus present in Adélie penguin was gained, and hence to estimate the ages of these loci. Distributions of estimated ages for loci in intergenic, intronic, exonic, and regulatory sequence are shown in Supplementary Fig. 4. Supplementary Fig. 5 shows the numbers of inferred gains and losses of microsatellites on each edge of the subtree, scaled according to both the length of the edge and the amount of sequence that can be aligned to the chicken genome. The total numbers of extant microsatellite loci whose origins were inferred to pre-date selected ancestral nodes are given in Supplementary Table 5.

The relative densities of microsatellite loci (including both pure and impure microsatellites) in different types of sequence for different age brackets are shown in Fig. 1. While the older age brackets contain fewer loci overall, those loci are much more likely to be found in regulatory or coding sequences. The percentage of loci found in regulatory or coding sequence for each bracket is shown in Supplementary Table 6. Microsatellite loci in regulatory or coding sequences thus appear to be conserved over longer periods on average than those in intergenic sequence or introns. This suggests that they are maintained by selection, be it directly for the presence of a microsatellite or for the surrounding sequence. Total numbers of loci genotyped in the Adélie penguin samples for each age bracket are given in Supplementary Table 7, along with the percentages of loci at which we observe multiple genotypes, showing that these loci retain length variability in Adélie penguins.

**Fig. 1.**
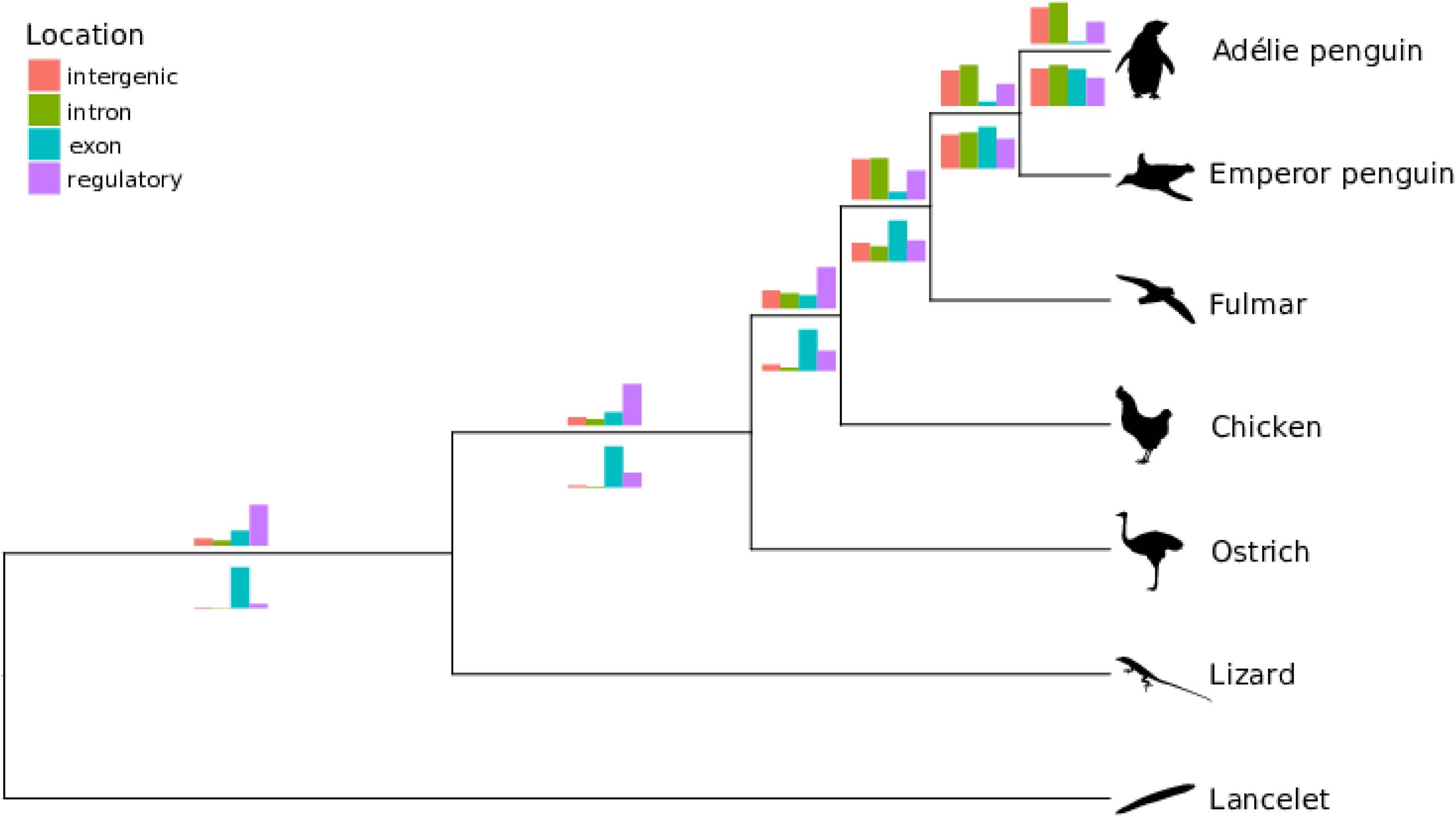
Relative densities of microsatellite loci in intergenic, intron, exon, and regulatory sequences in the Adélie penguin genome. Relative densities for loci inferred to have arisen on the branches shown on the tree. Those with periods two and three are displayed above and below the branches, respectively. Each plot shows relative rather than absolute densities, because the densities decrease rapidly with increasing locus age. Edge lengths are not drawn to scale.

### Microsatellite dynamics through deep time

To test whether the process of microsatellite mutation results in allele length distributions that are stationary over evolutionary time-scales (millions of years), we used BayesFactor as described above, replacing the sample age parameter with the locus age estimate. Bayes factors for all models tested are given in Supplementary Table 8. For all subsets of the data comprising pure and impure microsatellites of each period, the best-fitting model for allele length is dependent on motif, surrounding sequence type, locus age, and an interaction between surrounding sequence type and locus age. In all cases, the data provide very strong evidence for this model, being more likely under this model than under any other by a factor of at least 10^12^. We sampled from the posterior distribution of the full model for each subset to obtain posterior estimates of effect sizes, and these are shown in Supplementary Table 9. The effect of locus age is shown separately in Table 1. The effect sizes are very small (on the order of one nucleotide per hundred million years). For loci of periods 2 and 3, we also tested interactions between motif and locus age, and between motif and surrounding sequence type for subsets of our data, and found strong evidence for these interactions. We were unable to test these interactions for loci of longer periods because of the rapid increase in numbers of motifs as period increases.

**Table 1:** Posterior mean effect of locus age on length.

| Period | Pure | Impure |
| --- | --- | --- |
| 2 | 0.0051 [0.0049–0.0052] | 0.0100 [0.0095–0.0105] |
| 3 | 0.0056 [0.0055–0.0058] | 0.0140 [0.0136–0.0145] |
| 4 | 0.0038 [0.0034–0.0040] | 0.0125 [0.0113–0.0137] |
| 5 | 0.0058 [0.0048–0.0067] | 0.0113 [0.0097–0.0128] |
| 6 | 0.0010 [0.0006–0.0014] | 0.0120 [0.0111–0.0132] |

Mean inferred rate of increase in microsatellite length, in nucleotides per million years. Numbers in brackets represent 95% highest posterior density intervals.

Distributions of allele lengths for loci of different ages in different types of surrounding sequence are shown in Fig. 2. In agreement with the results of the linear mixed-model, a very slow increase in mean allele length over time can be seen for microsatellites in intron and intergenic sequence. Overall, pure di- and tetranucleotide loci in protein-coding sequence have significantly shorter mean allele lengths than those in non-coding sequence, while impure tri- and hexanucleotide microsatellites in protein-coding sequence have significantly longer mean allele lengths than those in non-coding sequence (see Table 2). It is likely that selection against frameshift mutations in coding sequence limits microsatellite expansion when the period is not a multiple of three^26^.

**Table 2:** Mean and standard error of allele lengths at microsatellite loci in different types of surrounding sequence.

| Period | Pure |  |  |  | Impure |  |  |  |
| --- | --- | --- | --- | --- | --- | --- | --- | --- |
|  | Intergenic | Intron | Exon | Regulatory | Intergenic | Intron | Exon | Regulatory |
| 2 | 13.06 | 12.93 | 11.79 | 12.79 | 20.53 | 20.11 | 19.66 | 21.03 |
|  | [0.0027] | [0.0039] | [0.0121] | [0.0100] | [0.0080] | [0.0129] | [0.0718] | [0.0303] |
| 3 | 15.99 | 15.70 | 16.06 | 16.03 | 24.02 | 23.68 | 26.73 | 22.79 |
|  | [0.0048] | [0.0071] | [0.0154] | [0.0158] | [0.0162] | [0.0302] | [0.0521] | [0.0402] |
| 4 | 16.40 | 16.03 | 14.83 | 15.61 | 24.59 | 24.12 | 24.45 | 23.15 |
|  | [0.0043] | [0.0058] | [0.0258] | [0.0103] | [0.0111] | [0.0174] | [0.1324] | [0.0360] |
| 5 | 18.93 | 18.35 | 17.35 | 17.70 | 27.79 | 26.84 | 24.44 | 25.86 |
|  | [0.0091] | [0.0127] | [0.0468] | [0.0226] | [0.0156] | [0.0244] | [0.1561] | [0.0524] |
| 6 | 18.44 | 18.02 | 17.78 | 17.95 | 27.73 | 26.74 | 29.71 | 26.75 |
|  | [0.0086] | [0.0124] | [0.0137] | [0.0280] | [0.0187] | [0.0274] | [0.1027] | [0.0870] |

**Fig. 2.**
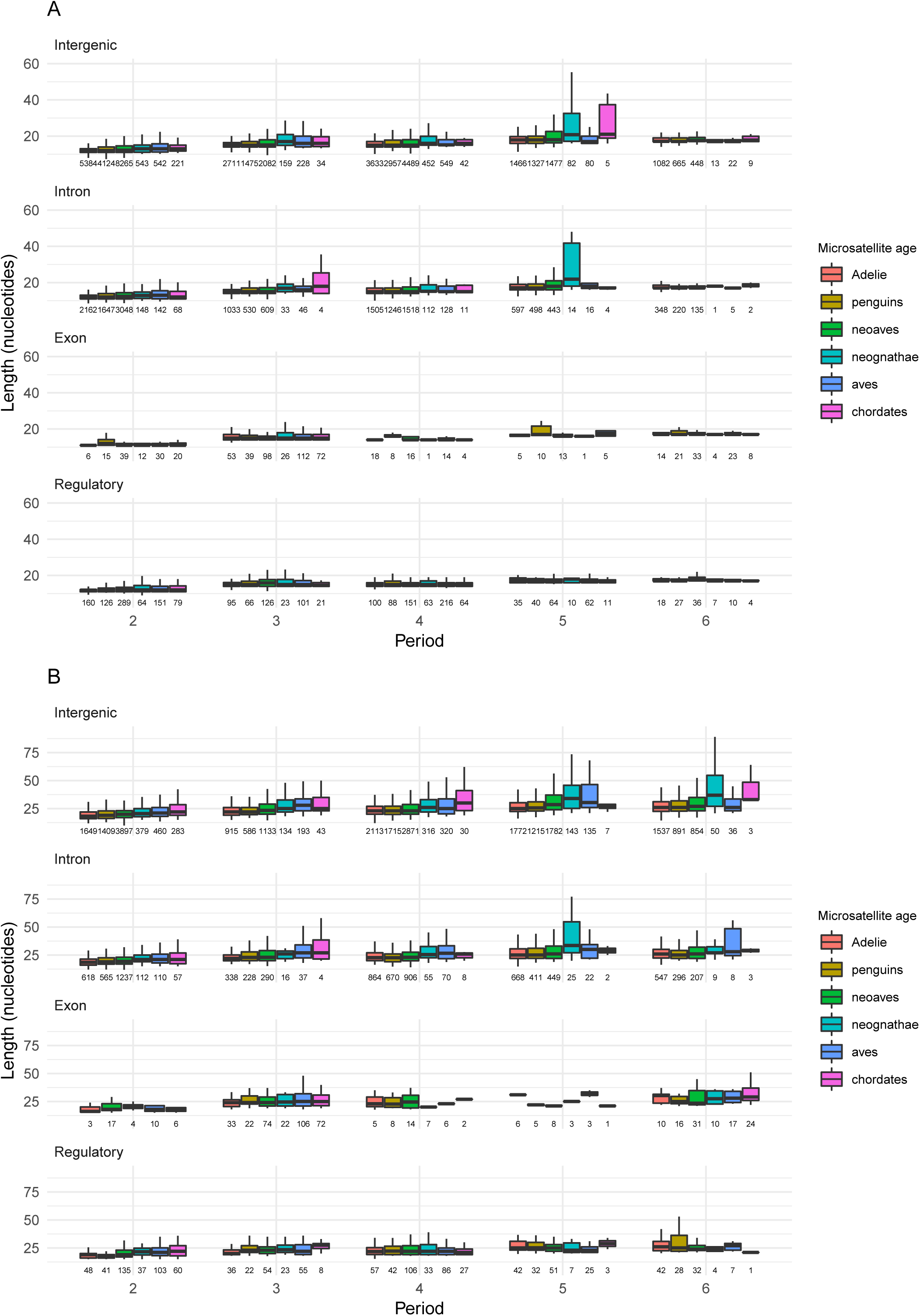
Distributions of allele lengths for loci of different ages in different types of surrounding sequence. Distributions of mean allele lengths (in nucleotides) of pure (A) and impure (B) microsatellite loci present in Adélie penguin and conserved across six age brackets, for loci with periods two to six in intergenic, intron, exon, and regulatory sequences. The six age brackets in each cluster correspond to loci that arose most recently on the branch leading to Adélie penguin; on the branch leading to penguins; within neoaves or on the branch leading to neoaves; on the branch leading to neognathae; on the branch leading to birds; outside sauria. (Note that no pure exonic microsatellites of period 5 are inferred to have arisen outside sauria.) Each box extends from the lower to upper quartiles of the length distribution, and the interior line indicates the median. The whiskers extend to the most extreme points within (1.5 × interquartile range) of the quartiles. Total numbers of loci are shown below each box.

## Discussion

To summarize our results, the mean allele length at any given microsatellite locus changes very little, on scales ranging from a few thousand to hundreds of millions of years, with estimated effect sizes on the order of one nucleotide per hundred million years. There is a gradual increase in allele length variation over time, as can be seen in Fig. 2. This suggests that the replication slippage process that generates length polymorphism is in a dynamic equilibrium, such that increases and decreases in length remain approximately balanced. These results are consistent with the findings of Sun *et al.*^18^ that longer alleles tend to decrease in length and shorter alleles tend to increase. We recommend that population geneticists and ecologists use models of microsatellite evolution that have stationary distributions, such as those of Kruglyak *et al.*^11^, Calabrese *et al.*^12^ or Amos *et al.*^13^, rather than those, such as the stepwise mutation model^27^, that allow allele lengths to drift upwards indefinitely.

We have also shown that microsatellites can persist, and remain variable, over very long periods of evolutionary time, with 257 extant microsatellite loci dating from before the origin of chordates, and 3,938 pre-dating the divergence of mammals and reptiles. Although we observe a slight decrease in heterozygosity with locus age (Supplementary Fig. 6), nevertheless, we observe multiple alleles in the Adélie samples for many ancient loci (Supplementary Table 7). The microsatellite loci that persist over very long periods are more often found in coding sequence and in regulatory regions. A disproportionate number of these variable ancient loci are trimer repeats located in protein-coding genes, which must code for a homopolymer run of amino acids. These trimer repeats in coding sequences make up only 0.55% of all loci that are variable in our Adélie samples, but 5.67% of variable loci that pre-date the divergence of extant birds, and 9.86% of those that pre-date the divergence of mammals and reptiles. It seems likely that selection is acting to maintain variability at these loci, which could act as mediators of rapid phenotypic change^2^.

A limitation of using short read data is that longer alleles are effectively censored from our data; however, as can be seen in Supplementary Fig. 1, the overwhelming majority of loci are much shorter (in the reference genome) than the read length. In addition, the reads from ancient Adélie samples are shorter than those from modern samples. This means that longer alleles are less likely to be genotyped in the ancient samples, and therefore we would expect this to give a signal for increasing allele length over time. However, we do not observe any such signal despite this potential bias, presumably because any such signal is swamped by the much larger number of shorter loci. As long-read sequencing becomes more common, and as methods for genotyping microsatellites in long-read data are developed, it may become feasible to verify our results for a more complete data set.

## Materials and Methods

### Contemporary Adélie penguin samples

Blood samples from Adélie penguins were collected from individuals at active breeding colonies, using methods as described in Millar *et al.*^28^, in six locations around Antarctica: Tongerson Island (AP samples) the Mawson region (B samples), Cape Adare (CA), Cape Bird (CB), Coulman Island (CI), and Inexpressible Island (II). Collection and sequencing information is given in Supplementary Table 10.

### Ancient Adélie penguin samples

Sub-fossil bones were collected in abandoned nests discovered along coastal ice-free areas both in the vicinity of presently occupied colonies and in relict colonies discovered in sites where penguins do not breed at present^29-31^ (Supplementary Table 11). Ornithogenic soils were stratigraphically excavated to find penguin bones and other remains as described previously^32,33^.

Radiocarbon AMS dates were supplied by NOSAMS, Woods Hole Oceanographic Institute, the New Zealand Institute of Geological and Nuclear Sciences, Lower Hutt, New Zealand, and Institut for Fysik og Astronomi, Aarhus Universitet, Denmark. Radiocarbon dates were calibrated with CALIB 7.1 (http://calib.qub.ac.uk/calib/)^34^ using the Marine Reservoir Correction Database 2013 and applying a delta-*R* of 791 ± 121 [^35^]. Mean ages and 2 delta standard deviation values were considered.

### Modern DNA extraction

For the 26 modern Adélie penguin samples, genomic libraries were prepared by first extracting DNA from Seutin-preserved blood or ethanol-preserved soft tissue samples. DNA was then purified using Qiagen DNEasy spin-columns according to the manufacturer’s protocol (Qiagen, Valencia, CA, USA) and eluted in 100 µL UltraPure™ water (Life Technologies, Grand Island, New York, USA).

### Ancient DNA extraction

All laboratory work with ancient Adélie penguin samples prior to PCR-amplification of genomic libraries (see below) was carried out in a physically isolated laboratory used only for ancient DNA work, following strict guidelines to minimize external contamination. Designated blank samples consisting originally of 200 µL digestion buffer were carried sequentially through all DNA extraction and library building procedures at a minimum ratio of one blank for every eight samples.

DNA was extracted from ancient bone or muscle tissue samples by first digesting ca. 0.1 g bone/tissue shavings in 200 µL digestion buffer (consisting of 180 µL of 0.5 M EDTA, 10 µL of 10% N-lauryl sarcosine, 10 µL of 20mg/mL proteinase K) for 12–18 hours at 55°C with rotational mixing (ca. 10 rpm). This was followed by 2–5 rounds of organic extraction with 1– 1.5 mL ultra-pure buffer-saturated phenol and one round of extraction with 1–1.5 mL chloroform (Sigma-Aldrich, St. Louis, MO, USA). Extracts were purified with Qiagen MinElute or PCR Purification columns using high concentration buffer PB or PE (10:1 buffer:sample volume ratio) to improve retention of small fragments, and 2× spin-through centrifugation for sample application and elution stages to further maximize yield. Final elutions were completed in a volume of 22 µL UltraPure™ water or NEB buffer EB.

### DNA library construction

Purified extracted DNA of modern samples was quantified with a Qubit 2.0 Fluorometer and dsDNA HS Assay kit (Life Technologies, Grand Island, NY, USA) and ca. 0.5–1.5 µg of DNA was sheared with a Covaris sonicator (Covaris, Woburn, MA, USA) to an average size of 300–600 bp (base pairs). Sheared extracts were adapter-ligated and enriched using the standard NEBNext or NEBNext Ultra protocol (catalog #E6040 and #E7370) and NEBNext multiplex Illumina primers (catalog #E7335) (NEB, Ipswich, MA, USA) in ½-size recommended reaction volumes for end-preparation, adapter ligation, and enrichment reactions. Enrichments were performed under recommended cycling conditions, with 10–14 cycles of enrichment for each sample and using Phusion High-Fidelity Master Mix (NEB catalog #M0531). Samples were submitted for 101 bp paired-end sequencing on an Illumina HiSeq2000 at BGI-Hong Kong, using one or ca. 1.33 lanes for each sample.

Genomic libraries for ancient samples were built following two strategies. Library building for all Holocene samples and initial attempts for two late Pleistocene samples (CB070121.08, CB070121.16) were completed based on Meyer *et al.*^36^ with minor adjustments. Based on low endogenous yields for the two late Pleistocene samples, a second attempt at library building was made for all three late Pleistocene samples (CB070121.08, CB070121.13, CB070121.16) following the NEBNext Ultra protocol, and using ½-size reactions for end-preparation, adapter ligation and enrichment reactions. For all ancient samples, enrichment reactions were completed by mixing ca. 11.5 µL of the heat-inactivated adapter-ligation reaction, 0.5 µL each of 25 µM NEBNext index and universal primers, and 12.5 µL 2× Phusion Hi-Fidelity Master Mix. Enrichment reactions were carried out under recommended cycling conditions, with 12–22 cycles of enrichment for each sample.

Finished ancient libraries were purified using Axygen MAG-PCR SPRI beads (Corning Life Sciences, Tewksbury, MA, USA) at a ratio of 0.7-1.1:1 Axygen:sample volume to minimize concentration of potential adapter dimers^37^ and quantified with a Qubit 2.0 Fluorometer. Libraries were submitted for 101bp single-end (SE) sequencing on an Illumina HiSeq2000 to either BGI-Hong Kong or the National High-Throughput Sequencing Center (University of Denmark, http://seqcenter.ku.dk/), using from between two and 10.5 lanes of sequencing for each sample with resultant genome-wide average sequencing depths of ca. 22× and 8× for modern and ancient samples, respectively (Supplementary Tables 10 and 11).

### Alignment

For all sequence pools, adapter sequences were trimmed from reads using Cutadapt^38^ v. 1.1 under default parameters. Low-quality reads were filtered with Trimmomatic^39^ v. 0.22, with minimum trailing and leading quality of 20, average quality over 20bp sliding windows of 20, and minimum lengths of 80bp for modern reads and 30bp for ancient reads. Trimmed and filtered Illumina reads for each Adélie penguin sample were mapped to the Adélie reference genome^22^ using Bowtie2^40^ with the ‘--very-sensitive’ preset option.

### Genomes

In this study we use the 48 avian genomes reported by Jarvis *et al.*^20^. We also use the pairwise alignments to the chicken genome that were used by Jarvis *et al.* in generating their whole-genome multiple alignment. This consists of a set of pairwise alignments for each species with each individual chromosome of the chicken genome as reference.

In addition, we use genomes of the fifteen non-avian species for which whole-genome alignments to the chicken galGal3 assembly are available from the UCSC genome browser. These are: human (hg19), chimpanzee (panTro3), orangutan (ponAbe2), mouse (mm9), rat (rn4), guinea pig (cavPor3), horse (equCab2), opossum (monDom5), platypus (ornAna1), lizard (anoCar2), frog (xenTro3), zebrafish (danRer4), fugu (fr2), lamprey (petMar1), and lancelet (braFlo1). All genomes used are listed in Supplementary Table 12.

### Microsatellite detection

Microsatellite loci were identified in all 63 genomes using Tandem Repeats Finder (TRF)^41^ with the following parameters: match weight 2; mismatch weight 7; indel weight 7; matching probability 80; indel probability 10; minimum alignment score 18; maximum period size 6. The results were then filtered using the alignment score thresholds shown in Supplementary Table 13, taken from Willems *et al.*^42^. This gave us five sets of microsatellites for each species: for dimer, trimer, tetramer, pentamer and hexamer repeats, with their respective score thresholds.

Microsatellite loci were compared against the annotations for all the avian genomes, to determine which loci fall within protein coding sequences or introns. Putative regulatory regions were identified by extracting the set of conserved nonexonic elements identified in the chicken genome by Lowe *et al.*^43^ from each of the avian genome alignments. All remaining sequences were assumed to be intergenic.

Microsatellites identified in the Adélie penguin reference genome using TRF were genotyped in the Adélie penguin samples using RepeatSeq^44^ (which requires a list of pre-identified loci and sequence reads as input), and the output formatted as tables for analysis in R^45^. Tables of genotype calls were imported into R and summary statistics calculated for each locus, including the mode, mean and standard deviation of the allele lengths observed in the samples, and the number of alleles observed. These were combined with the TRF output containing the motif, purity and nucleotide composition of the locus in the reference genome.

### Homology matching

First, for each species and period, we coded the microsatellite loci detected above as features in a general feature format file. Next, we used MafFilter^46^ to extract these features from the pairwise alignment between the species in question and each chicken chromosome, and output the coordinates that each feature aligns to in the chicken genome. A custom R script was used to produce a table matching each set of chicken coordinates to the corresponding microsatellite locus. Motifs were standardized by calculating the lexicographically minimal rotation to allow for loci to begin at different positions within a repeat unit (e.g., the motif TGA was standardized as ATG).

For each chicken chromosome and period, we combined the motif tables for all 63 species, and used a custom Java program to assign similarity scores to pairs of loci based on the distance between them (in terms of chicken coordinates) and the similarity of their motifs. Loci were scored if they were no more than 60 bp apart and their motifs differed by no more than one substitution. Testing different values of the length threshold showed that larger values did not increase the numbers of homologous loci detected. In addition, we manually checked a small sample of loci to verify that the loci detected were indeed homologous. Similarity was calculated as

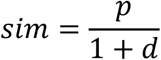

where *p* is the proportion of sites in the motif that are identical, and *d* is the distance in base pairs between the loci (zero if the loci overlap). We then used the Markov Cluster Algorithm (MCL)^47^ with the `--abc` input option and default settings to identify clusters of putatively homologous loci (across all 63 genomes). These clusters were converted into a matrix with the 48 species as columns and locations as rows, containing the motif for each species where a microsatellite is present. The matrix was also output as a presence/absence matrix, with ones where a microsatellite is present and zeroes otherwise.

To avoid any false negatives where a given region is not represented in the alignment for some species, we checked the local region of the alignment for any species missing from a given cluster, and recoded them as unknown (‘?’, as opposed to ‘0’ for absent) in the presence/absence matrix if the region was not covered in the alignment.

### Ancestral state reconstruction

We used the R package phangorn^48^ to perform ancestral state reconstruction on the timetree reported in Jarvis *et al.*^20^ using our presence/absence matrices. The maximum likelihood reconstructions available do not allow non-reversible models (i.e. the rates of gains and losses are assumed to be equal), so we used the “ACCTRAN” parsimony method. Numbers of gains and losses of homologous microsatellites inferred for each edge were then counted, ignoring any changes from a known state to unknown.

We also calculated the numbers of microsatellite losses required under a Dollo process, where any microsatellite locus only ever arises once, but may be lost in multiple lineages. However, the results were not appreciably different to those obtained under parsimony.

### Adélie locus age determination

Minimum ages were calculated for loci present in the Adélie penguin genome by using the ancestral state reconstruction results to identify the most recent gain of the locus on the path from the root to Adélie. This allows for loci being gained independently in different lineages, or lost and re-gained. These locus ages were combined with the genotype statistics calculated above, allowing us to examine the relationship between locus age, length, purity, and surrounding sequence type.

### Model fitting

The ‘generalTestBF’ function of the R package BayesFactor^21^ was used to fit generalized linear mixed models to the ancient and modern Adélie genotype data. A Bayes Factor (BF) is a measure that quantifies the evidence for a hypothesis compared to an alternative hypothesis given the data. The following thresholds have been suggested to quantify the evidence for one hypothesis over another as reported by BFs: BF < 3: insignificant, BF 3–20: positive, BF 20– 150: strong, BF > 150 very strong^49^.

We tested the dependence of microsatellite allele length on sample age, motif, sample, surrounding sequence type, and an interaction between sample age and surrounding sequence type for all loci genotyped in Adélie. For those loci for which we were able to estimate the age (i.e., those that were alignable to the chicken genome), we tested the dependence of allele length on estimated locus age, motif, sample, surrounding sequence type, and an interaction between locus age and surrounding sequence type. In both cases, the sample was treated as a random effect, and all other variables as fixed effects. Sample age and Locus age variables were centred. Models were fit separately for pure and impure microsatellite loci of each period (2 to 5). To obtain estimates of effect sizes, we used the ‘posterior’ function of BayesFactor to generate samples from the posterior distributions of the full models. We also tested interactions between motif and locus age, and between motif and surrounding sequence type for subsets of our data for loci of periods 2 and 3.

Our workflow for detecting homologous microsatellite loci and estimating their ages, starting from genome sequences and pairwise alignments, is given in Supplementary Fig. 7.

## Data and code availability

The datasets generated and analysed during the current study, and the code used for analysis, are available in the Dryad repository, https://doi.org/10.5061/dryad.7gt3rg2. The Adélie penguin sequence read data have been deposited with links to BioProject accession number PRJNA210803 in the NCBI BioProject database (https://www.ncbi.nlm.nih.gov/bioproject/).

## Acknowledgments

We thank John Macdonald and Peter Ritchie for assistance with collection of contemporary Adélie penguin samples.

## Author contributions

B.J.M., B.R.H., C.D.M. and D.M.L. conceived and, together with M.A.C., designed the study. D.M.L., C.D.M., and B.R.H. acquired funding. C.B. & M.C.S conducted geomorphologic field survey and discovered relict penguin colonies, sampled and dated in collaboration with M.P. and C.D.M. M.P., C.D.M. and D.M.L participated in collection of contemporary Adélie penguin samples. M.P. carried out DNA library construction. R.L. and G.Z. provided genome alignments. B.J.M., M.A.C. and B.R.H. analyzed the data. B.J.M., M.A.C., M.P., B.R.H. and D.M.L. wrote and revised the manuscript, with contributions from the other authors.

## References

1 Kashi, Y. & King, D. G. Simple sequence repeats as advantageous mutators in evolution. Trends Genet. 22, 253–259, doi: 10.1016/j.tig.2006.03.005 (2006).

2 Gemayel, R., Vinces, M. D., Legendre, M. & Verstrepen, K. J. Variable tandem repeats accelerate evolution of coding and regulatory sequences. Annu. Rev. Genet. 44, 445–477, doi: 10.1146/annurev-genet-072610-155046 (2010).

3 Mirkin, S. M. Expandable DNA repeats and human disease. Nature 447, 932–940, doi: 10.1038/nature05977 (2007).

4 Schlötterer, C. The evolution of molecular markers—just a matter of fashion? Nat. Rev. Genet. 5, 63–69, doi: 10.1038/nrg1249 (2004).

5 Ellegren, H. Microsatellite mutations in the germline: Implications for evolutionary inference. Trends Genet. 16, 551–558, doi: 10.1016/S0168-9525(00)02139-9 (2000).

6 Levinson, G. & Gutman, G. A. Slipped-strand mispairing: A major mechanism for DNA sequence evolution. Mol. Biol. Evol. 4, 203–221 (1987).

7 Bhargava, A. & Fuentes, F. F. Mutational dynamics of microsatellites. Mol. Biotechnol. 44, 250–266, doi: 10.1007/s12033-009-9230-4 (2010).

8 Schlötterer, C. Evolutionary dynamics of microsatellite DNA. Chromosoma 109, 365–371, doi: 10.1007/s004120000089 (2000).

9 Kelkar, Y. D., Eckert, K. A., Chiaromonte, F. & Makova, K. D. A matter of life or death: How microsatellites emerge in and vanish from the human genome. Genome Res. 21, 2038–2048, doi: 10.1101/gr.122937.111 (2011).

10 Buschiazzo, E. & Gemmell, N. J. The rise, fall and renaissance of microsatellites in eukaryotic genomes. Bioessays 28, 1040–1050, doi: 10.1002/bies.20470 (2006).

11 Kruglyak, S., Durrett, R. T., Schug, M. D. & Aquadro, C. F. Equilibrium distributions of microsatellite repeat length resulting from a balance between slippage events and point mutations. Proc. Natl. Acad. Sci. USA 95, 10774–10778, doi: 10.1073/pnas.95.18.10774 (1998).

12 Calabrese, P. P., Durret, R. T. & Aquadro, C. F. Dynamics of microsatellite divergence under stepwise mutation and proportional slippage/point mutation models. Genetics 159, 839–852, doi: melanogaster species complex drosophila-melanogaster saccharomyces-cerevisiae genetic distances range constraints point mutations tandem repeat allele size loci population (2001).

13 Amos, W., Kosanović, D. & Eriksson, A. Inter-allelic interactions play a major role in microsatellite evolution. Proc. R. Soc. Lond., Ser. B: Biol. Sci. 282, 20152125, doi: 10.1098/rspb.2015.2125 (2015).

14 Weber, J. L. & Wong, C. Mutation of human short tandem repeats. Hum. Mol. Genet. 2, 1123–1128 (1993).

15 Rose, O. & Falush, D. A threshold size for microsatellite expansion. Mol. Biol. Evol. 15, 613–615, doi: 10.1093/oxfordjournals.molbev.a025964 (1998).

16 Xu, X., Peng, M., Fang, Z. & Xu, X. The direction of microsatellite mutations is dependent upon allele length. Nat. Genet. 24, 396–399, doi: 10.1038/74238 (2000).

17 Huang, Q.-Y. et al. Mutation patterns at dinucleotide microsatellite loci in humans. Am. J. Hum. Genet. 70, 625–634, doi: 10.1086/338997 (2002).

18 Sun, J. X. et al. A direct characterization of human mutation based on microsatellites. Nat. Genet. 44, 1161–1165, doi: 10.1038/ng.2398 (2012).

19 Garza, J. C., Slatkin, M. & Freimer, N. B. Microsatellite allele frequencies in humans and chimpanzees, with implications for constraints on allele size. Mol. Biol. Evol. 12, 594–603 (1995).

20 Jarvis, E. D. et al. Whole-genome analyses resolve early branches in the tree of life of modern birds. Science 346, 1320–1331, doi: 10.1126/science.1253451 (2014).

21 Morey, R. D. & Rouder, J. N. Package “BayesFactor”. (2015). <https://cran.r->project.org/web/packages/BayesFactor/BayesFactor.pdf>.

22 Zhang, G. et al. Comparative genomics reveals insights into avian genome evolution and adaptation. Science 346, 1311–1320, doi: 10.1126/science.1251385 (2014).

23 Hedges, S. B., Marin, J., Suleski, M., Paymer, M. & Kumar, S. Tree of life reveals clock-like speciation and diversification. Mol. Biol. Evol. 32, 835–845, doi: 10.1093/molbev/msv037 (2015).

24 Prum, R. O. et al. A comprehensive phylogeny of birds (Aves) using targeted next-generation DNA sequencing. Nature, 3–11, doi: 10.1038/nature15697 (2015).

25 Suh, A., Smeds, L. & Ellegren, H. The dynamics of incomplete lineage sorting across the ancient adaptive radiation of neoavian birds. PLoS Biol. 13, e1002224, doi: 10.1371/journal.pbio.1002224 (2015).

26 Metzgar, D., Bytof, J. & Wills, C. Selection against frameshift mutations limits microsatellite expansion in coding DNA. Genome Res. 10, 72–80, doi: 10.1101/gr.10.1.72 (2000).

27 Ohta, T. & Kimura, M. A model of mutation appropriate to estimate the number of electrophoretically detectable alleles in a finite population. Genet. Res. 22, 201–204, doi: 10.1017/S0016672300012994 (1973).

28 Millar, C. D. et al. Mutation and evolutionary rates in Adélie penguins from the Antarctic. PLoS Genet. 4, e1000209, doi: 10.1371/journal.pgen.1000209 (2008).

29 Baroni, C. & Orombelli, G. Abandoned penguin rookeries as Holocene paleoclimatic indicators in Antarctica. Geology 22, 23–26, doi: 10.1130/0091-7613(1994)022<0023:APRAHP>2.3.CO;2 (1994).

30 Baroni, C. in Treatise on Geomorphology Vol. 8 (eds J. F. Schroeder, R. Giardino, & J. Harbor) 430–459 (Elsevier, 2013).

31 Lorenzini, S. et al. Adélie penguin dietary remains reveal Holocene environmental changes in the western Ross Sea (Antarctica). Palaeogeogr., Palaeoclimatol., Palaeoecol. 395, 21–28, doi: 10.1016/j.palaeo.2013.12.014 (2014).

32 Lambert, D. M. et al. Rates of evolution in ancient DNA from Adélie penguins. Science 295, 2270–2273, doi: 10.1126/science.1068105 (2002).

33 Ritchie, P. A., Millar, C. D., Gibb, G. C., Baroni, C. & Lambert, D. M. Ancient DNA enables timing of the Pleistocene origin and Holocene expansion of two Adélie penguin lineages in Antarctica. Mol. Biol. Evol. 21, 240–248, doi: 10.1093/molbev/msh012 (2004).

34 Reimer, P. J. et al. IntCal13 and Marine13 radiocarbon age calibration curves 0–50,000 years cal BP. Radiocarbon 55, 1869–1887, doi: 10.2458/azu_js_rc.55.16947 (2013).

35 Hall, B. L., Henderson, G. M., Baroni, C. & Kellogg, T. B. Constant Holocene Southern-Ocean 14C reservoir ages and ice-shelf flow rates. Earth Planet. Sci. Lett. 296, 115–123, doi: 10.1016/j.epsl.2010.04.054 (2010).

36 Meyer, M. & Kircher, M. Illumina sequencing library preparation for highly multiplexed target capture and sequencing. Cold Spring Harbor Protocols 5, pdb–prot5448, doi: 10.1101/pdb.prot5448 (2010).

37 Quail, M. A., Swerdlow, H. & Turner, D. J. Improved protocols for the Illumina genome analyzer sequencing system. Curr. Protoc. Hum. Genet. 62, 18.12.11-18.12.27, doi: 10.1002/0471142905.hg1802s62 (2009).

38 Martin, M. Cutadapt removes adapter sequences from high-throughput sequencing reads. EMBnet.journal 17, 10–12, doi: 10.14806/ej.17.1.200 (2011).

39 Bolger, A. M., Lohse, M. & Usadel, B. Trimmomatic: A flexible trimmer for Illumina sequence data. Bioinformatics 30, 2114–2120, doi: 10.1093/bioinformatics/btu170 (2014).

40 Langmead, B. & Salzberg, S. L. Fast gapped-read alignment with Bowtie 2. Nat. Methods 9, 357–359, doi: 10.1038/nmeth.1923 (2012).

41 Benson, G. Tandem repeats finder: A program to analyze DNA sequences. Nucleic Acids Res. 27, 573–580, doi: 10.1093/nar/27.2.573 (1999).

42 Willems, T. F., Gymrek, M., Highnam, G., Mittelman, D. & Erlich, Y. The landscape of human STR variation. Genome Res., 1894–1904, doi: 10.1101/gr.177774.114 (2014).

43 Lowe, C. B., Clarke, J. A., Baker, A. J., Haussler, D. & Edwards, S. V. Feather development genes and associated regulatory innovation predate the origin of Dinosauria. Mol. Biol. Evol. 32, 23–28, doi: 10.1093/molbev/msu309 (2015).

44 Highnam, G. et al. Accurate human microsatellite genotypes from high-throughput resequencing data using informed error profiles. Nucleic Acids Res. 41, e32, doi: 10.1093/nar/gks981 (2013).

45 R: A Language and Environment for Statistical Computing (R Foundation for Statistical Computing, Vienna, Austria, 2011).

46 Dutheil, J. Y., Gaillard, S. & Stukenbrock, E. H. MafFilter: A highly flexible and extensible multiple genome alignment files processor. BMC Genomics 15, 53, doi: 10.1186/1471-2164-15-53 (2014).

47 van Dongen, S. Graph clustering by flow simulation PhD thesis, University of Utrecht, (2000).

48 Schliep, K. P. phangorn: Phylogenetic analysis in R. Bioinformatics (Oxford, England) 27, 592–593, doi: 10.1093/bioinformatics/btq706 (2011).

49 Kass, R. E. & Raftery, A. E. Bayes Factors. Journal of the American Statistical Association 90, 773–795, doi: 10.1080/01621459.1995.10476572 (1995).

