## Supplementary materials for "Ancient and modern genomes reveal microsatellites maintain a dynamic equilibrium through deep time"

**This PDF file includes:**

Supplementary Figs. 1 to 7

Supplementary Tables 1 to 13

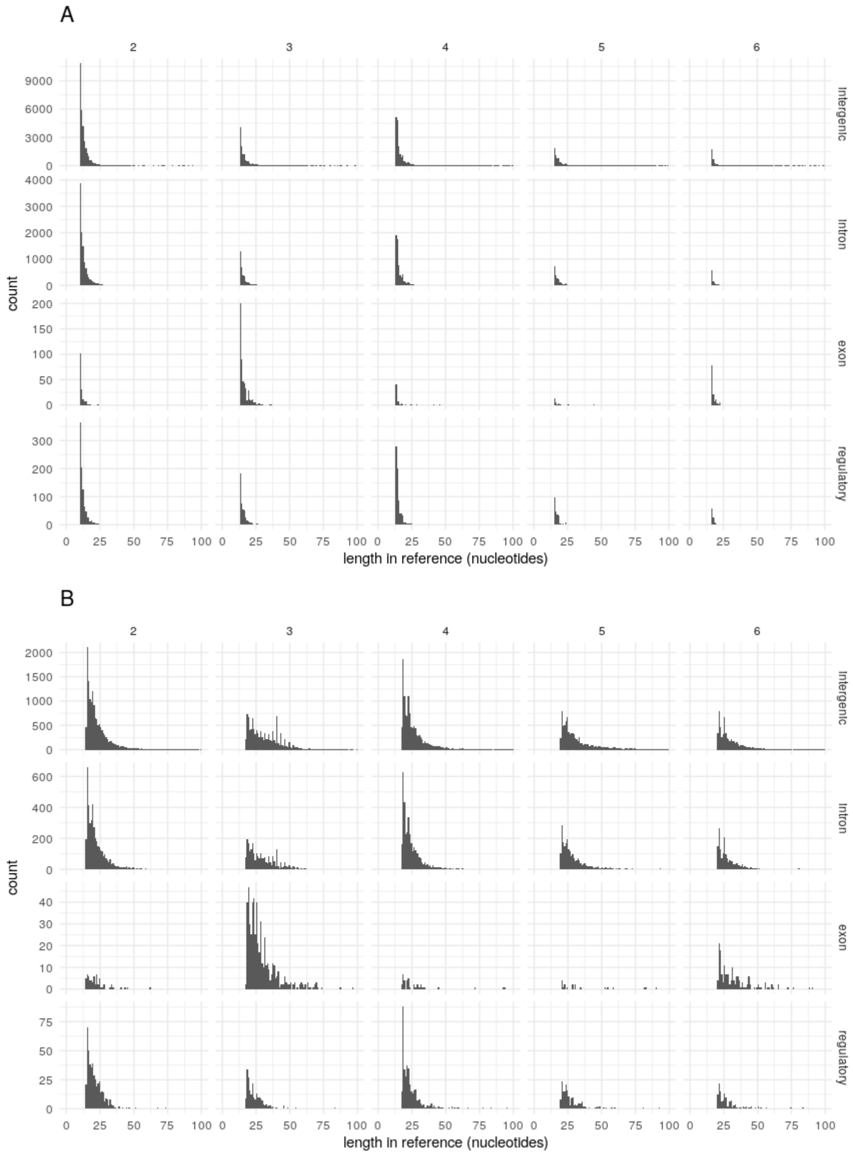


**Supplementary Fig. 1**

**Locus length distributions.** The distributions of microsatellite locus lengths in the Adélie penguin reference genome, for microsatellites of periods two to six, in intergenic, intronic, exonic, and regulatory sequence. Pure (A) and impure (B) loci are shown separately because the alignment score threshold effectively imposes different minimum observable lengths depending on the purity of the allele seen in the reference genome. For clarity, loci of length >100 bp are not shown, although small numbers of loci up to 6164 bp in length were detected.

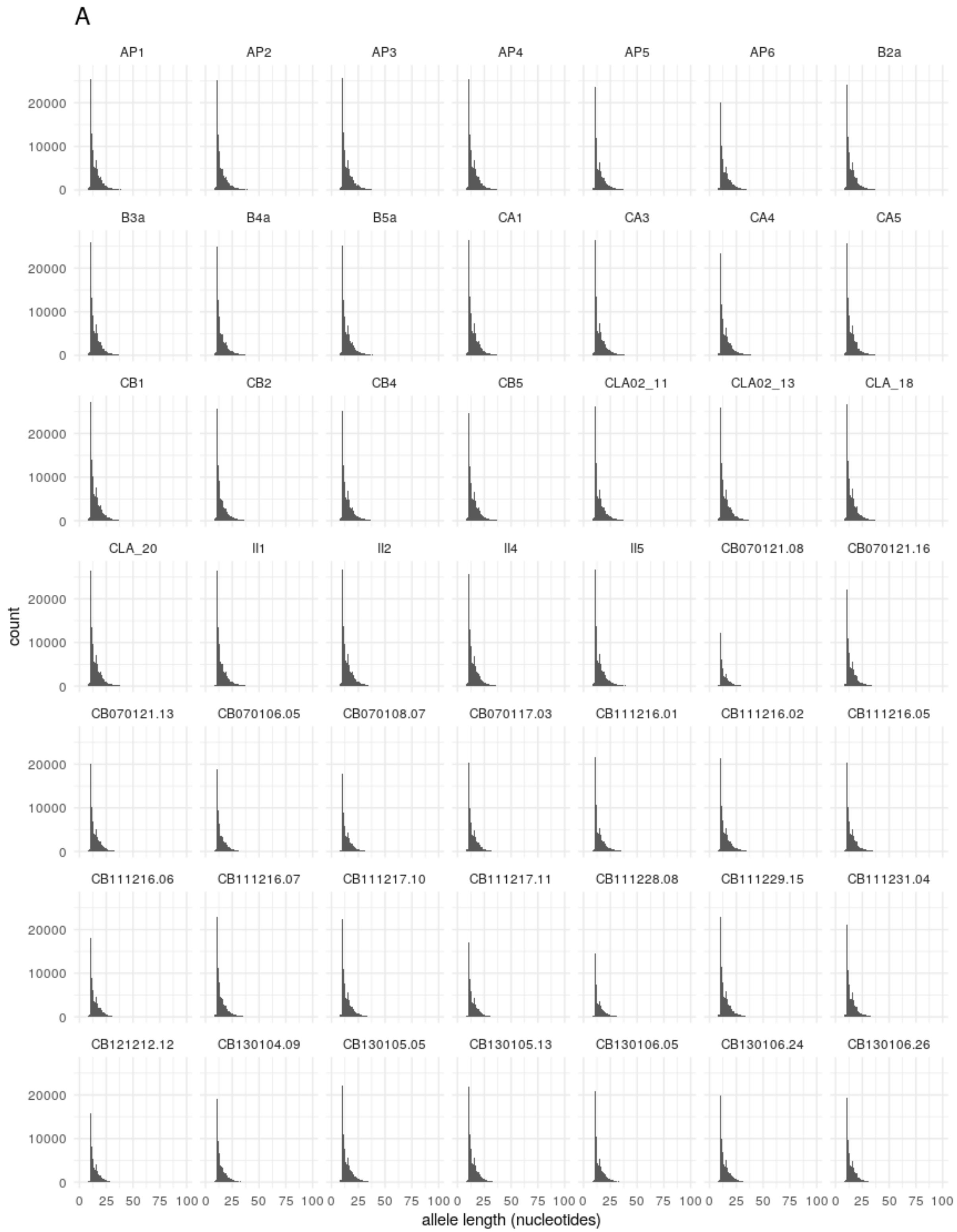
A
AP1
AP2
AP3
AP4
AP5
AP6
B2a
B3a
B4a
B5a
CA1
CA3
CA4
CA5
CB1
CB2
CB4
CB5
CLA02_11
CLA02_13
CLA_18
CLA_20
Il1
Il2
Il4
Il5
CB070121.08
CB070121.16
CB070121.13
CB070106.05
CB070108.07
CB070117.03
CB111216.01
CB111216.02
CB111216.05
CB111216.06
CB111216.07
CB111217.10
CB111217.11
CB111228.08
CB111229.15
CB111231.04
CB121212.12
CB130104.09
CB130105.05
CB130105.13
CB130106.05
CB130106.24
CB130106.26
count
0
10000
20000
0 25 50 75 100
allele length (nucleotides)

B

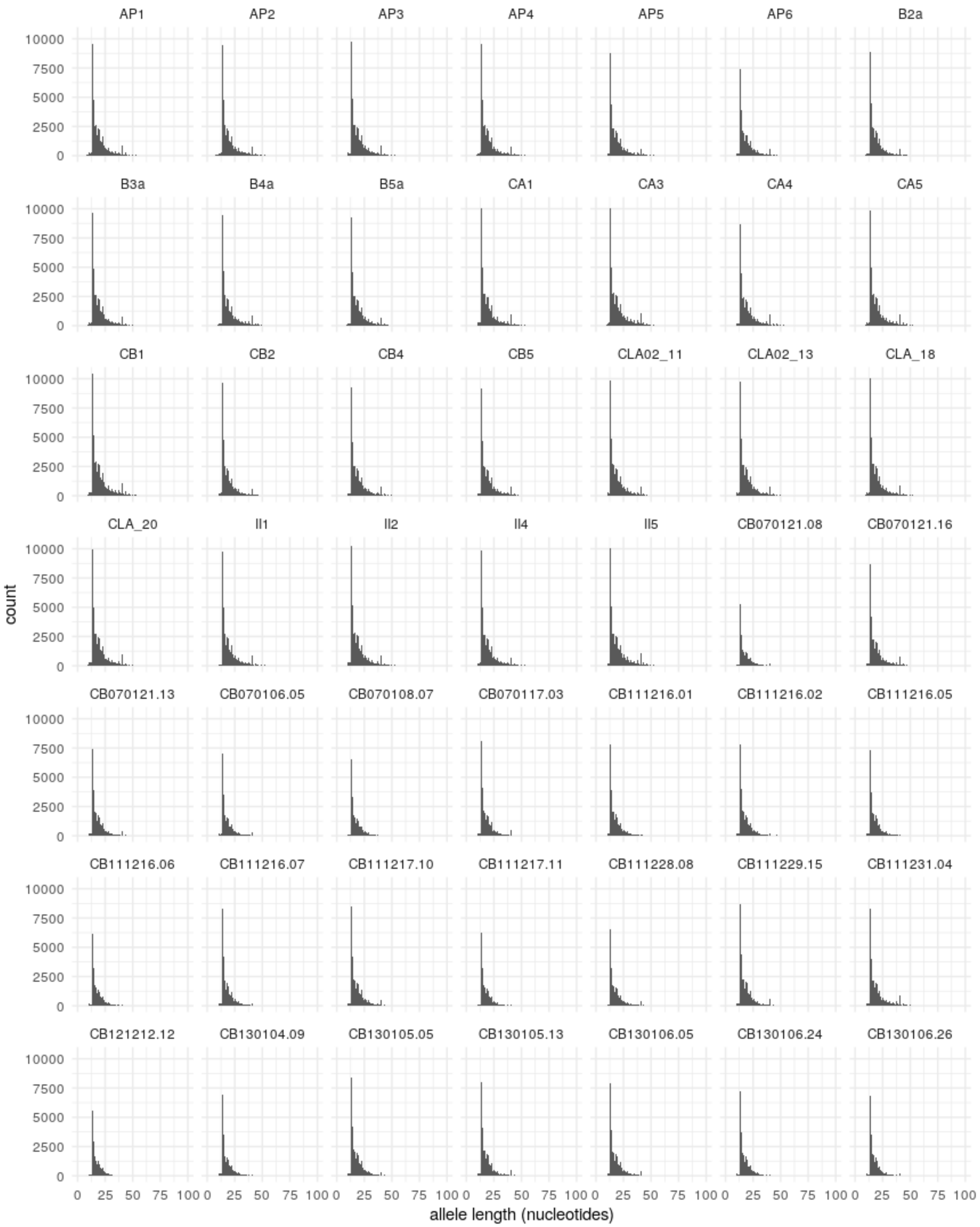
AP1
AP2
AP3
AP4
AP5
AP6
B2a
B3a
B4a
B5a
CA1
CA3
CA4
CA5
CB1
CB2
CB4
CB5
CLA02_11
CLA02_13
CLA_18
CLA_20
II1
II2
II4
II5
CB070121.08
CB070121.16
CB070121.13
CB070106.05
CB070108.07
CB070117.03
CB111216.01
CB111216.02
CB111216.05
CB111216.06
CB111216.07
CB111217.10
CB111217.11
CB111228.08
CB111229.15
CB111231.04
CB121212.12
CB130104.09
CB130105.05
CB130105.13
CB130106.05
CB130106.24
CB130106.26
count
0
2500
5000
7500
10000
0
25
50
75
100
allele length (nucleotides)

C

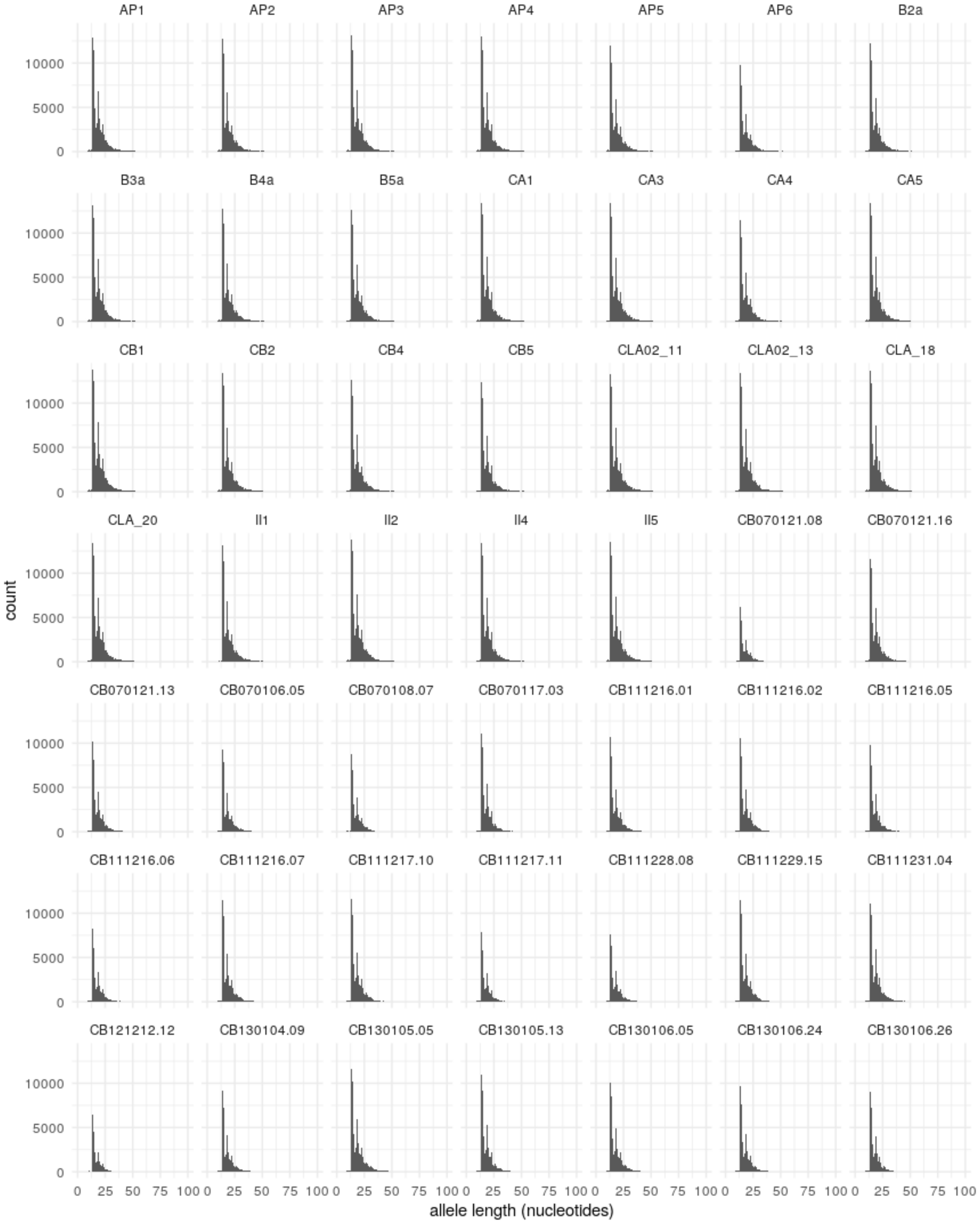
AP1
AP2
AP3
AP4
AP5
AP6
B2a
B3a
B4a
B5a
CA1
CA3
CA4
CA5
CB1
CB2
CB4
CB5
CLA02_11
CLA02_13
CLA_18
CLA_20
II1
II2
II4
II5
CB070121.08
CB070121.16
CB070121.13
CB070106.05
CB070108.07
CB070117.03
CB111216.01
CB111216.02
CB111216.05
CB111216.06
CB111216.07
CB111217.10
CB111217.11
CB111228.08
CB111229.15
CB111231.04
CB121212.12
CB130104.09
CB130105.05
CB130105.13
CB130106.05
CB130106.24
CB130106.26
count
0
5000
10000
0
25
50
75
100
allele length (nucleotides)

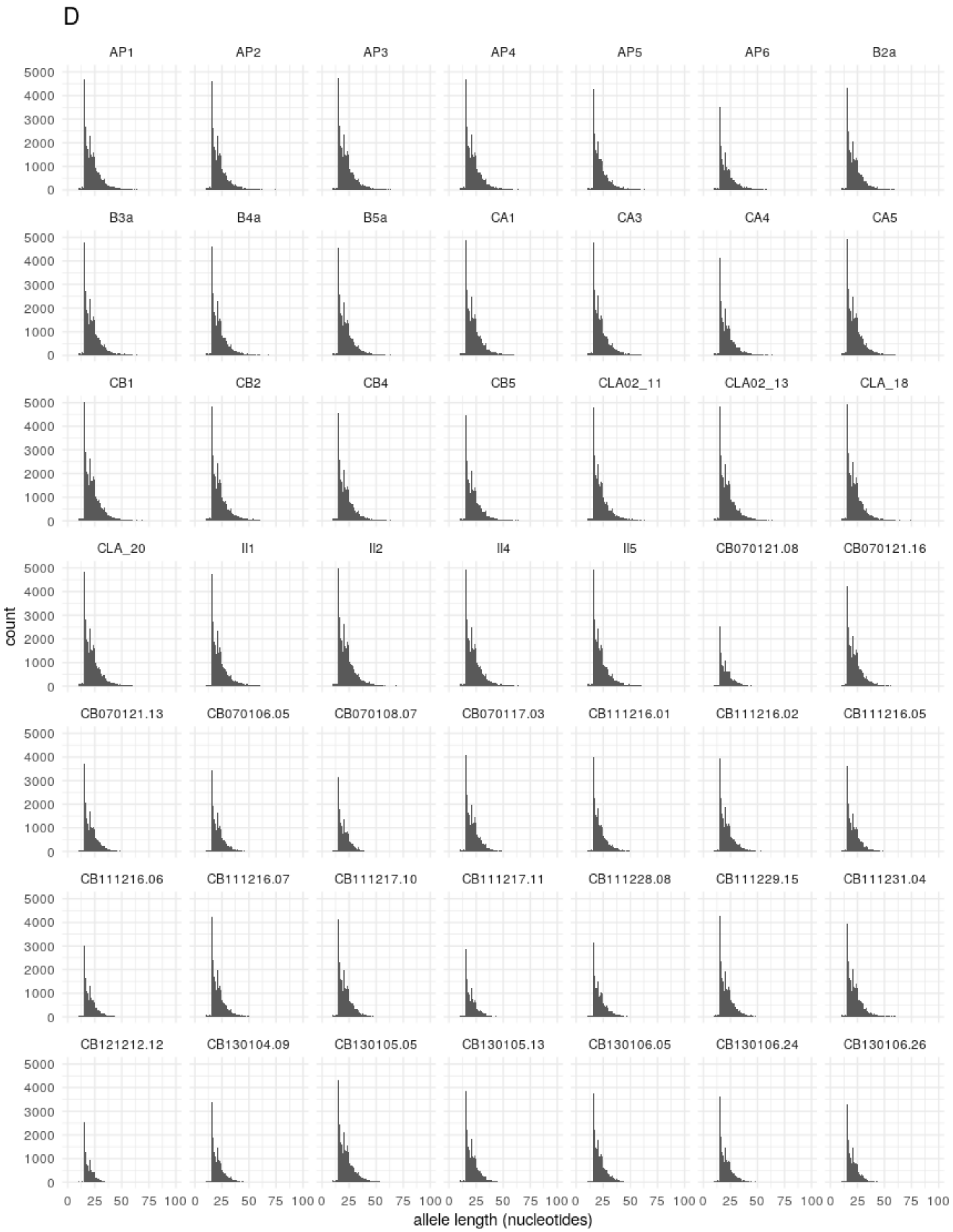
D
AP1
AP2
AP3
AP4
AP5
AP6
B2a
B3a
B4a
B5a
CA1
CA3
CA4
CA5
CB1
CB2
CB4
CB5
CLA02_11
CLA02_13
CLA_18
CLA_20
II1
II2
II4
II5
CB070121.08
CB070121.16
CB070121.13
CB070106.05
CB070108.07
CB070117.03
CB111216.01
CB111216.02
CB111216.05
CB111216.06
CB111216.07
CB111217.10
CB111217.11
CB111228.08
CB111229.15
CB111231.04
CB121212.12
CB130104.09
CB130105.05
CB130105.13
CB130106.05
CB130106.24
CB130106.26
count
0
1000
2000
3000
4000
5000
0 25 50 75 100
allele length (nucleotides)

E

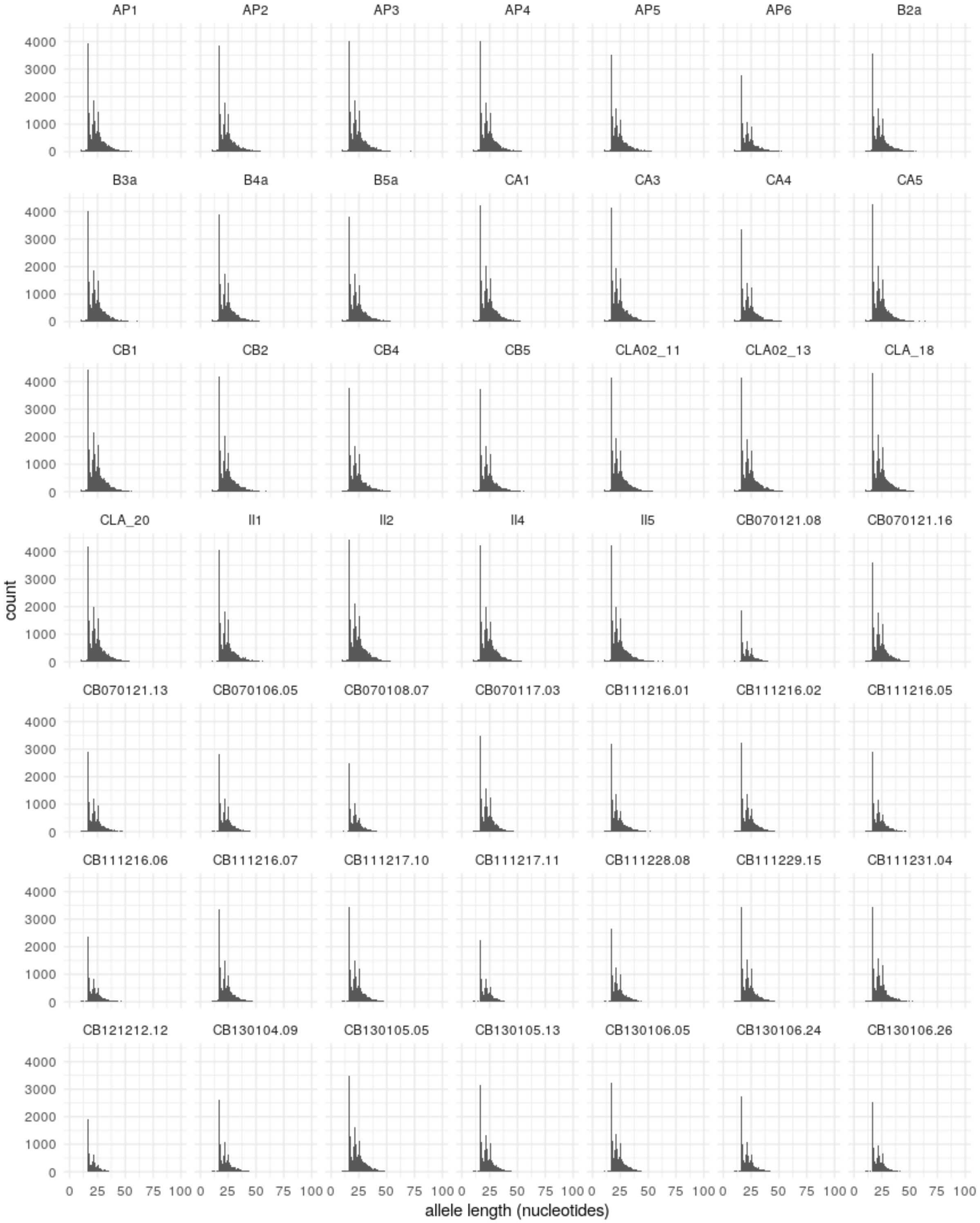
AP1
AP2
AP3
AP4
AP5
AP6
B2a
B3a
B4a
B5a
CA1
CA3
CA4
CA5
CB1
CB2
CB4
CB5
CLA02_11
CLA02_13
CLA_18
CLA_20
II1
II2
II4
II5
CB070121.08
CB070121.16
CB070121.13
CB070106.05
CB070108.07
CB070117.03
CB111216.01
CB111216.02
CB111216.05
CB111216.06
CB111216.07
CB111217.10
CB111217.11
CB111228.08
CB111229.15
CB111231.04
CB121212.12
CB130104.09
CB130105.05
CB130105.13
CB130106.05
CB130106.24
CB130106.26
count
0
1000
2000
3000
4000
0
25
50
75
100
allele length (nucleotides)

**Supplementary Fig. 2**

**Sample allele length distributions.** The distributions of microsatellite allele lengths genotyped in modern and ancient Adélie penguin samples, for microsatellites of periods (A) two to (E) six.

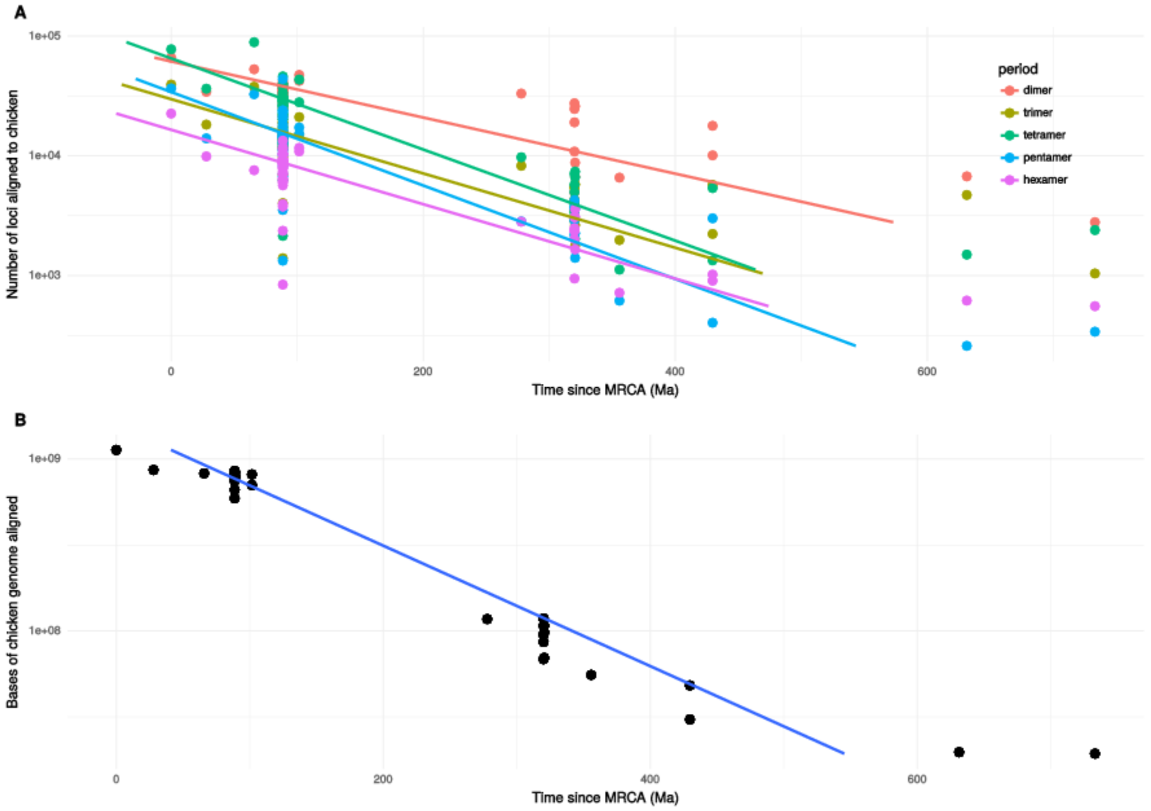


**Supplementary Fig. 3**

**Decrease in alignability over time.** Numbers of microsatellite loci in each species that can be aligned to the chicken genome (A) and overall sequence length aligned to chicken (B), plotted against the time since the most recent common ancestor of that species and chicken. Regression lines are calculated with time as the dependent variable, because the divergence times are estimates whereas the numbers of loci and overall sequence aligned are known precisely.

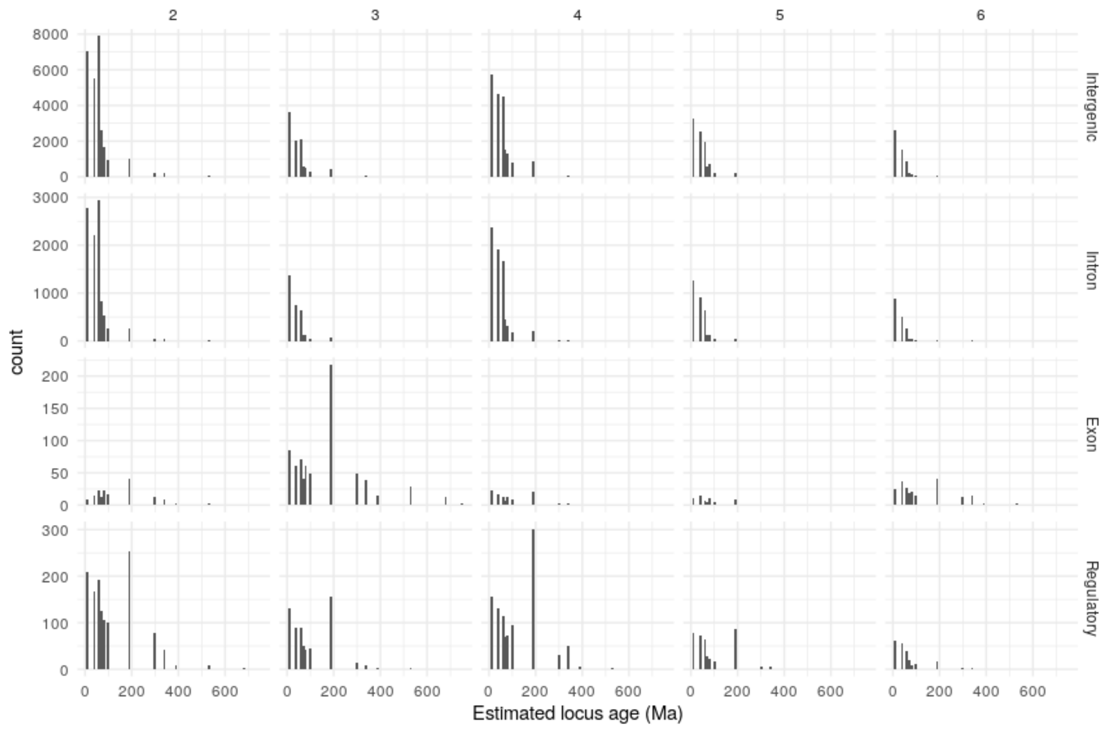


**Supplementary Fig. 4**

**Distribution of estimated locus ages.** The distributions of estimated ages of microsatellite loci of periods 2–6 in intergenic, intron, exon, and regulatory sequences in the Adélie penguin genome. Locus ages were estimated using the ancestral state reconstruction results to identify the most recent gain of the locus on the path from the root of the tree to Adélie.

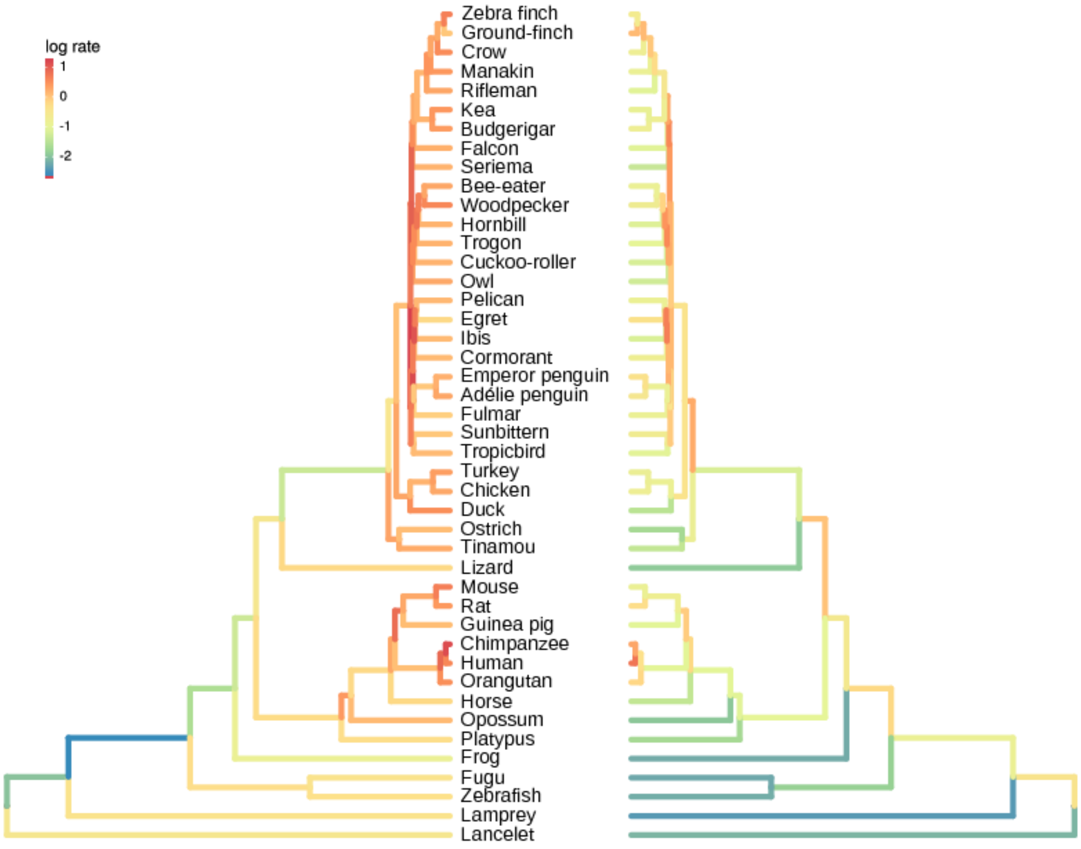


**Supplementary Fig. 5**

**Inferred rates of gain and loss of microsatellite loci.** The subtree used for ancestral state reconstruction, with edges colored according to the log of the number of microsatellite gains (left) and losses (right) per Mb of sequence alignable to the chicken genome per million years. Numbers of gains and losses are likely to be underestimated on longer edges due to multiple changes which are not detected by our ancestral state reconstruction.

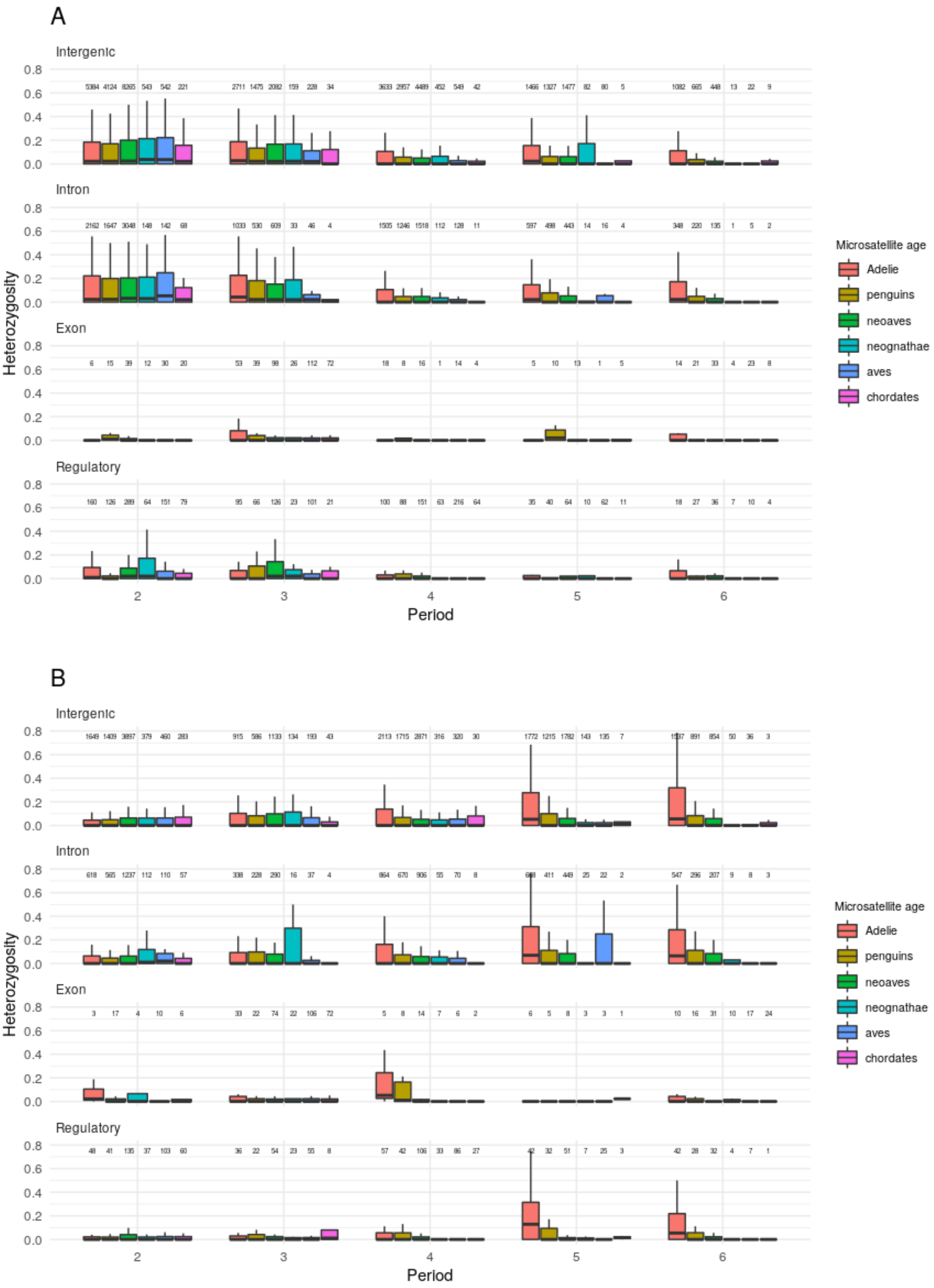
A
B
Intergenic
Intron
Exon
Regulatory
Heterozygosity
Period
Microsatellite age
Adelie
penguins
neoaves
neognathae
aves
chordates

**Supplementary Fig. 6**

**Microsatellite heterozygosity distributions.** Distributions of heterozygosity of pure (A) and impure (B) microsatellite loci present in Adélie penguin and conserved across six age brackets, for loci with periods two to six in intergenic, intron, exon, and regulatory sequences. The six age brackets in each cluster correspond, from left to right, to loci that arose most recently on the branch leading to Adélie penguin; on the branch leading to penguins; within neoaves or on the branch leading to neoaves; on the branch leading to neognathae; on the branch leading to birds; outside sauria. Each box extends from the lower to upper quartiles of the heterozygosity distribution, and the interior line indicates the median. The whiskers extend to the most extreme points within (1.5 × interquartile range) of the quartiles. Total numbers of loci are shown above each box.

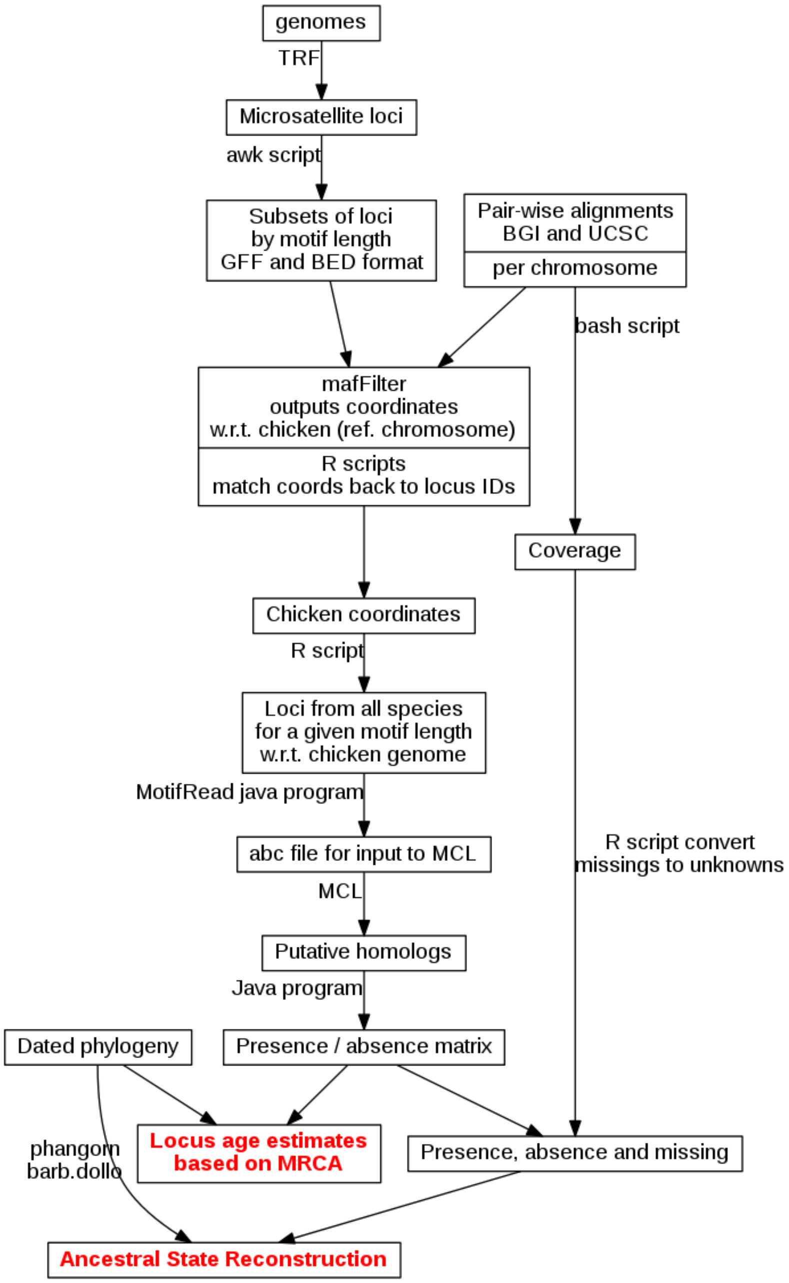


**Supplementary Fig. 7**

**Workflow diagram.** Our workflow for detecting homologous microsatellite loci and estimating their ages, starting from genome sequences and pairwise alignments.

**Supplementary Table 1: Comparison of different models with sample age as a variable to explain allele lengths of microsatellite loci**

| | Model | Period | | | | |
|---|---|---|---|---|---|---|
| | | 2 | 3 | 4 | 5 | 6 |
| Pure | sample + motif + location | 1 | 1 | 1 | 1 | 1 |
| | sample + motif + sample age + location | 0.0625 ±2.7% | 0.0536 ±1.84% | 0.0645 ±1.38% | 0.0711 ±2.87% | 0.0588 ±2.43% |
| | sample + motif + sample age + location + sample age * location | 2.95e-07 ±6.98% | 2.37e-06 ±2.14% | 6.70e-07 ±1.39% | 9.45e-07 ±2.26% | 9.36e-08 ±1.87% |
| | sample + motif | 7.41e-912 ±2.1% | 5.98e-1940 ±0.73% | 4.85e-1441 ±0.91% | 3.67e-767 ±0.74% | 4.02e-161 ±1.05% |
| | sample + motif + sample age | 4.93e-913 ±3.26% | 2.53e-693 ±1.51% | 3.21e-1442 ±1.29% | 2.53e-768 ±1.45% | 2.36e-162 ±1.5% |
| | sample + location | 2.36e-9825 ±2% | 2.02e-5597 ±1.76% | 3.22e-24549 ±1.02% | 1.83e-11911 ±1.09% | 1.63e-4623 ±1.19% |
| | sample + sample age + location | 1.36e-9826 ±3.23% | 1.09e-5598 ±1.76% | 2.11e-24550 ±1.53% | 1.26e-11912 ±1.94% | 8.45e-4625 ±1.54% |

| | | | | | | |
|---|---|---|---|---|---|---|
| | sample + sample age + location + sample age * location | 1.05e-9832 ±2.08% | 6.79e-5603 ±1.28% | 1.93e-24554 ±1.3% | 1.12e-11916 ±3.27% | 1.26e-4630 ±1.75% |
| | sample | 4.86e-10770 ±1.91% | 2.86e-6233 ±1.02% | 4.70e-26125 ±0.88% | 1.37e-12673 ±0.7% | 3.34e-4885 ±1.05% |
| | sample + sample age | 2.63e-10771 ±2.14% | 1.54e-6234 ±1.61% | 7.93e-26126 ±57.34% | 9.69e-12675 ±1.71% | 1.75e-4886 ±1.5% |
| Impure | sample + motif + location | 1 | 1 | 1 | 1 | 1 |
| | sample + motif + sample age + location | 0.0926 ±2.27% | 0.0691 ±1.68% | 0.135 ±1.73% | 0.102 ±2.66% | 0.103 ±1.92% |
| | sample + motif + sample age + location + sample age * location | 1.08e-06 ±2.86% | 4.86e-06 ±1.34% | 1.98e-04 ±1.44% | 1.91e-05 ±1.56% | 1.81e-06 ±1.06% |
| | sample + motif | 1.49e-447 ±1.26% | 5.98e-1940 ±0.73% | 2.76e-504 ±1.06% | 4.51e-434 ±1.26% | 1.36e-313 ±0.59% |
| | sample + motif + sample age | 1.41e-448 ±2.17% | 4.06e-1941 ±3.11% | 3.75e-505 ±2.02% | 4.72e-435 ±5.57% | 1.34e-314 ±1.33% |
| | sample + location | 2.34e-965 ±2.03% | 1.56e-7655 ±0.99% | 2.84e-16888 ±1.8% | 3.43e-12624 ±1.47% | 1.10e-14790 ±0.86% |
| | sample + sample age + location | 1.82e-966 ±1.7% | 1.77e-7656 ±1.63% | 3.65e-16889 ±2.03% | 3.46e-12625 ±4.9% | 1.44e-14791 ±12.82% |

| | | | | | | |
|---|---|---|---|---|---|---|
| | sample + sample age + location + sample age * location | 1.24e-971 ±1.44% | 1.02e-7662 ±1.4% | 4.74e-16891 ±1.46% | 1.31e-12628 ±1.69% | 1.73e-14796 ±14.13% |
| | sample | 8.69e-1420 ±1.08% | 6.28e-8813 ±0.8% | 4.74e-17497 ±1.05% | 2.31e-13327 ±1.33% | 1.27e-15600 ±0.58% |
| | sample + sample age | 7.24e-1421 ±3.13% | 6.77e-8814 ±1.43% | 6.10e-17498 ±1.39% | 2.31e-13328 ±1.69% | 1.39e-15601 ±1.01% |

Bayes factors for generalized linear mixed models in which allele length is treated as dependent on different combinations of motif, surrounding sequence type (exon, intron, regulatory, or intergenic), sample age, and interaction between surrounding sequence type and sample age. The sample, i.e., the particular Adélie genome, is treated as a random effect, and is present in all models. Models were fit separately for pure and impure microsatellites of each period. Bayes factors are relative to the best model (in all cases that in which allele length is dependent on the motif and surrounding sequence type).

**Supplementary Table 2: Posterior estimates of effect sizes in models including sample age as a variable**

| | | Parameter | Period | | | | |
|---|---|---|---|---|---|---|---|
| | | | 2 | 3 | 4 | 5 | 6 |
| Pure | | mu | 12.4227<br>[12.2811, 12.5736] | 16.1689<br>[16.0002, 16.3388] | 15.8932<br>[15.7177, 16.0760] | 18.9176<br>[18.6679, 19.1473] | 18.1558<br>[17.9944, 18.3212] |
| | | Sample age | -0.0050<br>[-0.0224, 0.0111] | -0.0036<br>[-0.0236, 0.0143] | -0.0061<br>[-0.0267, 0.0164] | -0.0059<br>[-0.0376, 0.0186] | -0.0057<br>[-0.0242, 0.0137] |
| | Location | Intergenic | 0.4438<br>[0.4283, 0.4598] | 0.2126<br>[0.1976, 0.2279] | 0.6196<br>[0.5912, 0.6487] | 0.6878<br>[0.6321, 0.7436] | 0.3625<br>[0.3312, 0.3930] |
| | | Intron | 0.2413<br>[0.2238, 0.2560] | -0.2838<br>[-0.3009, -0.2666] | 0.1202<br>[0.0915, 0.1518] | -0.1220<br>[-0.1789, -0.0656] | -0.0051<br>[-0.0417, 0.0302] |
| | | Exon | -0.7557<br>[-0.7961, -0.7150] | 0.1977<br>[0.1715, 0.2267] | -0.5398<br>[-0.6217, -0.4585] | -0.2648<br>[-0.4225, -0.1078] | -0.2402<br>[-0.2960, -0.1804] |
| | | Regulatory | 0.0706<br>[0.0474, 0.0924] | -0.1264<br>[-0.1558, -0.0917] | -0.2000<br>[-0.2358, -0.1617] | -0.3010<br>[-0.3753, -0.2272] | -0.1172<br>[-0.1882, -0.0559] |
| | Locus age * location | Intergenic | -0.0026<br>[-0.0044, -0.0008] | -0.0031<br>[-0.0050, -0.0014] | -0.0031<br>[-0.0065, 0.0008] | -0.0064<br>[-0.0127, 0.0004] | -0.0017<br>[-0.0055, 0.0019] |
| | | Intron | -0.0013<br>[-0.0033, 0.0007] | -0.0010<br>[-0.0032, 0.0013] | -0.0013<br>[-0.0048, 0.0026] | -0.0019<br>[-0.0095, 0.0045] | 0.0005<br>[-0.0041, 0.0045] |
| | | Exon | 0.0034<br>[-0.0019, 0.0080] | 0.0036<br>[0.0004, 0.0071] | -0.0011<br>[-0.0084, 0.0125] | 0.0079<br>[-0.0125, 0.0257] | 0.0032<br>[-0.0034, 0.0102] |
| | | Regulatory | 0.0006<br>[-0.0021, 0.0034] | 0.0005<br>[-0.0032, 0.0043] | 0.0033<br>[-0.0014, 0.0078] | -0.0004<br>[-0.0093, 0.0088] | -0.0019<br>[-0.0102, 0.0059] |
| Impure | | mu | 20.1424<br>[19.8505, 20.4129] | 26.8126<br>[26.1998, 27.3722] | 24.2881<br>[23.8806, 24.7417] | 28.3972<br>[27.9825, 28.8900] | 29.0784<br>[28.6993, 29.5254] |
| | | Sample age | -0.0224<br>[-0.0540, 0.0100] | -0.0170<br>[-0.0656, 0.0291] | -0.0277<br>[-0.0770, 0.0226] | -0.0213<br>[-0.0756, 0.0294] | -0.0257<br>[-0.0744, 0.0191] |
| | Location | Intergenic | -0.1199<br>[-0.1633, -0.0725] | 0.3386<br>[0.2922, 0.3892] | 0.6787<br>[0.6008, 0.7508] | 1.1114<br>[0.9960, 1.2482] | 0.3180<br>[0.2563, 0.3711] |
| | | Intron | -0.7473<br>[-0.7956, -0.6970] | -0.8378<br>[-0.8963, -0.7795] | -0.1154<br>[-0.1878, -0.0293] | 0.0434<br>[-0.0921, 0.1652] | -0.6507<br>[-0.7128, -0.5772] |

| | | | | | | | |
|---|---|---|---|---|---|---|---|
| | | Exon | 1.0625<br>[0.9407, 1.1851] | 3.3447<br>[3.2695, 3.4168] | -0.0092<br>[-0.2016, 0.2289] | -0.7002<br>[-1.0401, -0.3680] | 1.2848<br>[1.1577, 1.4012] |
| | | Regulatory | -0.1953<br>[-0.2558, -0.1303] | -2.8456<br>[-2.9649, -2.7493] | -0.5724<br>[-0.6749, -0.4680] | -0.4547<br>[-0.6193, -0.2918] | -0.9521<br>[-1.0912, -0.8170] |
| | Locus age * location | Intergenic | 0.0029<br>[-0.0030, 0.0085] | -0.0001<br>[-0.0060, 0.0055] | -0.0062<br>[-0.0162, 0.0023] | -0.0071<br>[-0.0227, 0.0058] | 0.0002<br>[-0.0068, 0.0068] |
| | | Intron | 0.0025<br>[-0.0034, 0.0088] | 0.0019<br>[-0.0045, 0.0093] | 0.0023<br>[-0.0076, 0.0121] | 0.0035<br>[-0.0124, 0.0174] | 0.0073<br>[-0.0006, 0.0145] |
| | | Exon | -0.0186<br>[-0.0330, -0.0027] | -0.0148<br>[-0.0239, -0.0067] | -0.0068<br>[-0.0316, 0.0217] | -0.0058<br>[-0.0480, 0.0348] | -0.0146<br>[-0.0277, -0.0015] |
| | | Regulatory | 0.0132<br>[0.0054, 0.0213] | 0.0130<br>[-0.0000, 0.0270] | 0.0107<br>[-0.0028, 0.0227] | 0.0094<br>[-0.0092, 0.0292] | 0.0070<br>[-0.0081, 0.0237] |

Means and 95% highest posterior density intervals of parameters sampled from the posterior distribution of a generalized linear mixed model in which allele length is treated as dependent on sample, motif, surrounding sequence type (exon, intron, regulatory, or intergenic), sample age, and interaction between surrounding sequence type and sample age. Units for allele length and sample age are nucleotides and thousands of years respectively. The effects of sample age and of the interaction term are therefore in nucleotides per thousand years.

**Supplementary Table 3: Numbers of microsatellite loci in reference genomes**

| Species | Period | | | | | Total | Genome size (Gb) | msats/Mb |
|---|---|---|---|---|---|---|---|---|
| | 2 | 3 | 4 | 5 | 6 | | | |
| Zebra finch | 67,108 | 33,531 | 50,898 | 33,063 | 23,511 | 208,111 | 1.256 | 165.6 |
| Medium ground-finch | 59,915 | 3,003 | 4,001 | 3,118 | 2,274 | 72,311 | 1.073 | 67.4 |
| American crow | 54,965 | 25,552 | 40,659 | 22,296 | 16,679 | 160,151 | 1.095 | 146.2 |
| Golden-collared manakin | 68,560 | 26,756 | 50,974 | 25,054 | 17,694 | 189,038 | 1.160 | 162.9 |
| Rifleman | 54,301 | 19,289 | 32,742 | 17,737 | 12,780 | 136,849 | 1.045 | 131.0 |
| Budgerigar | 55,005 | 21,767 | 39,580 | 18,668 | 13,524 | 148,544 | 1.133 | 131.1 |
| Kea | 51,577 | 20,220 | 41,873 | 22,483 | 14,886 | 151,039 | 1.062 | 142.2 |
| Peregrine falcon | 68,732 | 29,262 | 47,427 | 24,996 | 18,271 | 188,688 | 1.174 | 160.7 |
| Red-legged seriema | 54,052 | 21,610 | 40,156 | 23,444 | 13,870 | 153,132 | 1.146 | 133.6 |
| Carmine bee-eater | 50,662 | 24,213 | 49,302 | 28,382 | 14,932 | 167,491 | 1.067 | 157.0 |
| Downy woodpecker | 72,765 | 36,622 | 65,780 | 104,075 | 21,227 | 300,469 | 1.175 | 255.8 |

| | | | | | | | | |
|---|---|---|---|---|---|---|---|---|
| Rhinoceros hornbill | 50,975 | 24,443 | 36,300 | 20,582 | 13,021 | 145,321 | 1.106 | 131.4 |
| Bar-tailed trogon | 50,699 | 20,110 | 37,559 | 19,468 | 13,499 | 141,335 | 1.086 | 130.2 |
| Cuckoo-roller | 56,897 | 22,889 | 41,955 | 24,026 | 14,568 | 160,335 | 1.148 | 139.6 |
| Speckled mousebird | 52,208 | 18,342 | 31,832 | 18,512 | 9,582 | 130,476 | 1.086 | 120.1 |
| Barn owl | 59,714 | 23,887 | 55,381 | 35,030 | 19,119 | 193,131 | 1.139 | 169.6 |
| Bald eagle | 68,017 | 33,478 | 48,796 | 26,674 | 21,026 | 197,991 | 1.280 | 154.6 |
| White-tailed eagle | 56,513 | 11,618 | 18,900 | 9,814 | 5,980 | 102,825 | 1.145 | 89.8 |
| Turkey vulture | 58,661 | 25,051 | 43,490 | 21,604 | 13,926 | 162,732 | 1.180 | 137.9 |
| Dalmatian pelican | 57,783 | 32,483 | 42,116 | 23,063 | 14,120 | 169,565 | 1.169 | 145.1 |
| Little egret | 58,039 | 7,863 | 11,206 | 6,308 | 4,543 | 87,959 | 1.212 | 72.6 |
| Crested ibis | 64,942 | 29,446 | 50,507 | 28,755 | 17,462 | 191,112 | 1.240 | 154.1 |
| Great cormorant | 50,939 | 23,088 | 38,805 | 21,485 | 13,889 | 148,206 | 1.155 | 128.3 |
| Northern fulmar | 56,579 | 16,620 | 29,084 | 16,196 | 10,260 | 128,739 | 1.153 | 111.6 |
| Adelie penguin | 63,965 | 32,338 | 44,004 | 23,252 | 14,415 | 177,974 | 1.251 | 142.3 |
| Emperor penguin | 65,435 | 29,780 | 45,366 | 24,195 | 14,967 | 179,743 | 1.283 | 140.1 |

| | | | | | | | | |
|---|---|---|---|---|---|---|---|---|
| Red-throated loon | 51,932 | 18,162 | 35,238 | 17,614 | 11,244 | 134,190 | 1.145 | 117.2 |
| White-tailed tropicbird | 52,789 | 21,978 | 39,451 | 20,679 | 13,433 | 148,330 | 1.167 | 127.1 |
| Sunbittern | 47,957 | 10,906 | 25,164 | 10,934 | 6,427 | 101,388 | 1.099 | 92.3 |
| Killdeer | 59,920 | 26,159 | 46,762 | 28,132 | 18,175 | 179,148 | 1.225 | 146.2 |
| Grey-crowned crane | 59,469 | 27,358 | 45,743 | 23,963 | 14,732 | 171,265 | 1.139 | 150.4 |
| Hoatzin | 47,276 | 19,697 | 32,620 | 16,719 | 10,718 | 127,030 | 1.209 | 105.1 |
| Annas hummingbird | 67,426 | 31,290 | 61,754 | 40,394 | 25,075 | 225,939 | 1.115 | 202.7 |
| Chimney swift | 65,958 | 27,967 | 59,951 | 28,586 | 16,713 | 199,175 | 1.125 | 177.0 |
| Chuck-wills widow | 71,985 | 34,254 | 58,368 | 33,811 | 20,699 | 219,117 | 1.145 | 191.3 |
| MacQueens bustard | 52,657 | 21,894 | 42,495 | 23,737 | 14,267 | 155,050 | 1.095 | 141.5 |
| Red-crested turaco | 56,111 | 24,580 | 41,745 | 23,438 | 14,618 | 160,492 | 1.176 | 136.5 |
| Common cuckoo | 50,230 | 19,818 | 34,987 | 19,051 | 12,787 | 136,873 | 1.159 | 118.1 |
| Brown mesite | 64,474 | 27,361 | 43,458 | 27,666 | 19,307 | 182,266 | 1.101 | 165.5 |
| Yellow-throated sandgrouse | 53,507 | 22,055 | 38,378 | 23,186 | 13,875 | 151,001 | 1.098 | 137.6 |
| Pigeon | 60,907 | 29,113 | 71,736 | 38,871 | 18,978 | 219,605 | 1.112 | 197.6 |

| | | | | | | | | |
|---|---|---|---|---|---|---|---|---|
| American flamingo | 56,389 | 23,127 | 42,734 | 22,441 | 12,357 | 157,048 | 1.145 | 137.2 |
| Great crested grebe | 60,441 | 24,447 | 58,076 | 33,472 | 18,118 | 194,554 | 1.151 | 169.0 |
| Chicken | 69,385 | 39,912 | 79,481 | 37,204 | 22,759 | 248,741 | 1.127 | 220.7 |
| Turkey | 54,788 | 29,378 | 62,609 | 27,683 | 20,253 | 194,711 | 1.080 | 180.3 |
| Peking duck | 86,595 | 55,394 | 126,395 | 54,582 | 32,206 | 355,172 | 1.105 | 321.4 |
| White-throated tinamou | 95,271 | 33,771 | 69,074 | 28,411 | 20,363 | 246,890 | 1.060 | 232.8 |
| Common ostrich | 91,024 | 23,128 | 51,094 | 23,401 | 18,668 | 207,315 | 1.228 | 168.8 |
| Carolina anole | 498,193 | 371,856 | 219,382 | 47,725 | 57,686 | 1,194,842 | 1.835 | 651.1 |
| Human | 438,720 | 126,489 | 355,182 | 141,729 | 118,234 | 1,180,354 | 3.200 | 368.9 |
| Chimpanzee | 870,509 | 124,795 | 352,679 | 146,303 | 116,222 | 1,610,508 | 3.374 | 477.3 |
| Orangutan | 905,652 | 138,158 | 364,823 | 156,997 | 119,917 | 1,685,547 | 3.516 | 479.4 |
| Mouse | 1,262,387 | 180,711 | 549,771 | 180,770 | 182,531 | 2,356,170 | 2.780 | 847.5 |
| Rat | 1,152,208 | 142,738 | 411,864 | 114,999 | 148,218 | 1,970,027 | 2.891 | 681.5 |
| Guinea pig | 652,417 | 79,607 | 229,405 | 127,172 | 92,974 | 1,181,575 | 2.778 | 425.4 |
| Horse | 673,239 | 66,320 | 161,405 | 48,549 | 46,250 | 995,763 | 2.534 | 392.9 |

| | | | | | | | | |
|---|---|---|---|---|---|---|---|---|
| Opossum | 1,164,701 | 157,487 | 398,520 | 125,825 | 149,329 | 1,995,862 | 3.678 | 542.7 |
| Platypus | 422,056 | 419,757 | 316,734 | 33,006 | 36,090 | 1,227,643 | 2.037 | 602.7 |
| Western clawed frog | 373,377 | 40,266 | 63,865 | 14,364 | 16,976 | 508,848 | 1.542 | 330.0 |
| Zebrafish | 927,344 | 190,668 | 384,057 | 97,635 | 56,469 | 1,656,173 | 1.810 | 914.9 |
| Fugu | 169,548 | 23,068 | 30,046 | 12,211 | 14,930 | 249,803 | 0.408 | 611.5 |
| Sea lamprey | 449,151 | 130,995 | 65,362 | 15,454 | 14,777 | 675,739 | 1.048 | 644.9 |
| Lancelet | 169,323 | 29,568 | 80,898 | 20,471 | 32,634 | 332,894 | 0.944 | 352.3 |

Numbers of loci of periods 2–6 detected using TRF are shown for each of the 63 genomes analysed. Genome sizes and overall number of microsatellite loci per megabase are also given.

**Supplementary Table 4: Numbers of microsatellites detected and aligned**

| | | | | **Number of loci in:** | | | | |
|---|---|---|---|---|---|---|---|---|
| **Period** | **Detected** | **Aligned** | **Loci** | **1 sp.** | **2 spp.** | **3 spp.** | **4–10 spp.** | **>10 spp.** |
| **2** | 13,034,324 | 1,921,221 | 835,504 | 584,701 | 109,769 | 39,967 | 66,198 | 34,869 |
| **3** | 3,427,493 | 771,350 | 414,191 | 322,316 | 44,228 | 14,890 | 23,135 | 9,622 |
| **4** | 6,189,999 | 1,496,806 | 787,409 | 586,985 | 99,769 | 35,768 | 46,229 | 18,658 |
| **5** | 2,529,465 | 779,421 | 512,227 | 418,217 | 52,007 | 17,109 | 19,238 | 5,656 |
| **6** | 1,949,034 | 441,006 | 330,452 | 286,760 | 26,377 | 7,266 | 8,105 | 1,944 |
| **Total** | 27,130,315 | 5,409,804 | 2,879,783 | 2,198,979 | 332,150 | 115,000 | 162,905 | 70,749 |

Total numbers of microsatellites of periods 2–6 detected across the 63-taxon tree, numbers alignable to the chicken genome, and numbers of loci in the chicken genome to which these were mapped. For those loci in the chicken genome, numbers with microsatellites present in one, two, three, four to ten, and eleven or more species.

**Supplementary Table 5: Total numbers of extant microsatellite loci inferred to pre-date selected ancestral nodes**

| Node | Period | | | | |
|---|---|---|---|---|---|
| | **2** | **3** | **4** | **5** | **6** |
| **Chordata** | 164 | 74 | 13 | 0 | 6 |
| **Vertebrata** | 470 | 195 | 16 | 2 | 8 |
| **Gnathostomata** | 1,098 | 376 | 128 | 11 | 23 |
| **Amniota** | 2,365 | 808 | 526 | 122 | 117 |
| **Sauria** | 2,918 | 1,173 | 728 | 177 | 183 |
| **Aves** | 6,826 | 3,155 | 3,613 | 969 | 550 |

**Supplementary Table 6: Percentages of microsatellite loci of different ages found in coding or regulatory sequence in the Adélie penguin genome**

| | Age bracket | Period 2 | Period 3 | Period 4 | Period 5 | Period 6 |
|---|---|---|---|---|---|---|
| **Pure** | Adélie | 2.2 | 3.8 | 2.2 | 1.9 | 2.2 |
| | penguins | 2.4 | 5.0 | 2.2 | 2.7 | 5.1 |
| | neoaves | 2.8 | 7.7 | 2.7 | 3.9 | 10.6 |
| | neognathae | 9.9 | 20.3 | 10.2 | 10.3 | 44.0 |
| | birds | 20.9 | 43.7 | 25.4 | 41.1 | 55.0 |
| | outside sauria | 25.5 | 71.0 | 56.2 | 55.0 | 52.2 |
| **Impure** | Adélie | 2.2 | 5.2 | 2.0 | 1.9 | 2.4 |
| | penguins | 2.0 | 5.1 | 2.1 | 2.2 | 3.6 |
| | neoaves | 2.9 | 8.3 | 3.1 | 2.6 | 5.6 |
| | neognathae | 7.7 | 23.1 | 9.7 | 5.6 | 19.2 |
| | birds | 16.5 | 41.2 | 19.1 | 15.1 | 35.3 |
| | outside sauria | 16.3 | 63.0 | 43.3 | 30.8 | 80.6 |

Age brackets correspond to loci that arose most recently on the branch leading to Adélie penguin; on the branch leading to penguins; within neoaves or on the branch leading to neoaves; on the branch leading to neognathae; on the branch leading to birds; outside sauria.

**Supplementary Table 7: Numbers of microsatellite loci of different ages genotyped in Adélie penguin samples, and percentages of loci at which multiple genotypes are observed**

| | | Period | | | | |
|---|---|---|---|---|---|---|
| | **Age bracket** | **2** | **3** | **4** | **5** | **6** |
| **Pure** | Adélie | 7514 (58.6%) | 3818 (61.2%) | 5198 (47.0%) | 2030 (56.4%) | 1414 (51.6%) |
| | penguins | 5745 (58.9%) | 2060 (56.9%) | 4250 (41.6%) | 1806 (45.6%) | 917 (39.5%) |
| | neoaves | 11333 (60.9%) | 2808 (57.4%) | 6100 (39.5%) | 1792 (45.4%) | 641 (32.9%) |
| | neognathae | 735 (62.3%) | 223 (60.1%) | 615 (38.2%) | 86 (55.8%) | 23 (4.3%) |
| | birds | 839 (59.6%) | 471 (46.9%) | 892 (29.6%) | 150 (24.7%) | 59 (8.5%) |
| | outside sauria | 376 (53.7%) | 125 (39.2%) | 121 (16.5%) | 20 (30.0%) | 23 (21.7%) |
| **Impure** | Adélie | 1985 (38.8%) | 1188 (47.3%) | 2888 (49.0%) | 2286 (65.8%) | 1966 (64.3%) |
| | penguins | 1796 (38.8%) | 790 (42.7%) | 2373 (43.2%) | 1546 (50.1%) | 1178 (46.9%) |
| | neoaves | 4865 (42.7%) | 1431 (43.0%) | 3727 (41.0%) | 1973 (48.5%) | 1021 (41.5%) |
| | neognathae | 471 (43.5%) | 174 (44.8%) | 367 (41.4%) | 132 (48.5%) | 62 (33.9%) |
| | birds | 619 (42.2%) | 365 (41.4%) | 443 (36.6%) | 135 (37.8%) | 59 (23.7%) |

| | | | | | | |
|---|---|---|---|---|---|---|
| | outside sauria | 352 (43.8%) | 119 (35.3%) | 57 (31.6%) | 11 (45.5%) | 29 (13.8%) |

Age brackets correspond to loci that arose most recently on the branch leading to Adélie penguin; on the branch leading to penguins; within neoaves or on the branch leading to neoaves; on the branch leading to neognathae; on the branch leading to birds; outside sauria. Numbers in parentheses represent the percentage of loci at which multiple alleles are observed in Adélie penguin samples.

**Supplementary Table 8: Comparison of different models with locus age as a variable to explain allele lengths of microsatellite loci**

| | Model | Period | | | | |
|---|---|---|---|---|---|---|
| | | 2 | 3 | 4 | 5 | 6 |
| Pure | sample + motif + locus age + location + locus age * location | 1 | 1 | 1 | 1 | 1 |
| | sample + motif + locus age + location | 6.13e-640 ±2.41% | 3.04e-1011 ±1.79% | 6.28e-543 ±1.43% | 7.89e-165 ±7.96% | 5.14e-13 ±1.78% |
| | sample + motif + locus age | 1.13e-2062 ±2.08% | 4.43e-1381 ±2.86% | 3.33e-1852 ±1.29% | 1.21e-759 ±5.42% | 1.37e-158 ±2.27% |
| | sample + motif + location | 1.06e-7088 ±1.85% | 7.27e-2527 ±1.24% | 5.05e-2208 ±1.1% | 4.31e-722 ±1.48% | 7.26e-27 ±1.06% |
| | sample + motif | 4.86e-7594 ±1.95% | 1.64e-2744 ±1.1% | 6.82e-3034 ±0.84% | 6.27e-1164 ±1.14% | 1.57e-157 ±0.88% |
| | sample + locus age + location + locus age * location | 2.14e-5198 ±1.89% | 1.04e-2151 ±1.33% | 6.18e-12275 ±1.91% | 8.39e-6605 ±2.05% | 3.76e-4442 ±1.18% |
| | sample + locus age + location | 1.86e-5868 ±2.53% | 6.80e-3258 ±3.25% | 5.40e-12799 ±4.12% | 1.66e-6815 ±30.86% | 1.20e-4456 ±3.84% |
| | sample + locus age | 7.52e-7456 ±2.52% | 1.34e-3597 ±1.48% | 2.11e-14218 ±1.47% | 2.17e-7532 ±2.03% | 4.79e-4657 ±1.22% |
| | sample + location | 2.99e-12627 ±1.82% | 4.17e-4853 ±1.55% | 3.53e-14106 ±1.73% | 3.99e-7477 ±1.24% | 9.75e-4469 ±1.05% |
| | sample | 9.52e-13193 ±1.7% | 5.36e-5065 ±1.09% | 2.12e-15043 ±0.81% | 2.44e-7939 ±1.13% | 7.24e-4655 ±0.88% |
| Impure | sample + motif + locus age + location + locus age * location | 1 | 1 | 1 | 1 | 1 |
| | sample + motif + locus age + location | 1.98e-227 ±2.69% | 3.95e-436 ±3.1% | 9.94e-412 ±2.83% | 3.42e-357 ±1.57% | 7.76e-198 ±1.76% |

| | | | | | | |
|---|---|---|---|---|---|---|
| | sample + motif + locus age | 2.22e-424 ±2.79% | 1.30e-758 ±1.81% | 5.74e-925 ±4.22% | 1.88e-724 ±4.34% | 1.76e-355 ±6.68% |
| | sample + motif + location | 2.99e-5960 ±1.7% | 1.90e-1908 ±1.67% | 1.80e-2173 ±1.34% | 2.16e-1583 ±6.2% | 1.48e-895 ±2.55% |
| | sample + motif | 7.90e-6198 ±1.32% | 1.09e-2950 ±1.55% | 1.62e-2401 ±1.13% | 7.34e-1832 ±0.7% | 5.69e-1003 ±1.15% |
| | sample + locus age + location + locus age * location | 1.66e-495 ±2.06% | 2.98e-604 ±4.15% | 9.66e-8069 ±1.52% | 6.05e-5421 ±1.18% | 2.77e-8272 ±1.69% |
| | sample + locus age + location | 1.41e-722 ±2.32% | 5.85e-1063 ±2.1% | 6.73e-8559 ±2.01% | 1.02e-5841 ±2.89% | 5.89e-8579 ±1.67% |
| | sample + locus age | 2.06e-936 ±2.5% | 9.37e-1369 ±1.91% | 3.07e-9144 ±1.99% | 3.43e-6344 ±2.64% | 8.13e-8721 ±1.55% |
| | sample + location | 2.88e-6523 ±1.4% | 1.01e-2613 ±1.67% | 1.01e-10312 ±1.28% | 3.82e-7685 ±0.88% | 1.28e-9419 ±1.32% |
| | sample | 2.28e-6763 ±1.22% | 1.40e-3669 ±1.52% | 4.79e-10579 ±1.12% | 3.11e-8031 ±0.68% | 9.27e-9731 ±1.14% |

Bayes factors for generalized linear mixed models in which allele length is treated as dependent on different combinations of motif, surrounding sequence type (exon, intron, regulatory, or intergenic), estimated microsatellite locus age, and interaction between surrounding sequence type and locus age. The sample, i.e., the particular Adélie genome, is treated as a random effect, and is present in all models. Models were fit separately for pure and impure microsatellites of each period. Bayes factors are relative to the best model (the full model in all cases).

**Supplementary Table 9: Posterior estimates of effect sizes in models including sample age as a variable**

| | | Parameter | Period | | | | |
|---|---|---|---|---|---|---|---|
| | | | 2 | 3 | 4 | 5 | 6 |
| Pure | | mu | 12.2241 [12.0956, 12.3547] | 15.8446 [15.6915, 16.0037] | 15.7361 [15.5697, 15.9071] | 18.1887 [17.9187, 18.3909] | 18.2412 [18.1013, 18.3836] |
| | | Sample age | 0.0051 [0.0049, 0.0052] | 0.0056 [0.0055, 0.0058] | 0.0037 [0.0034, 0.0040] | 0.0058 [0.0048, 0.0067] | 0.0010 [0.0006, 0.0014] |
| | Location | Intergenic | 0.4310 [0.4120, 0.4513] | 0.1107 [0.0951, 0.1268] | 0.5481 [0.5135, 0.5799] | 0.6422 [0.5813, 0.7052] | 0.3271 [0.2900, 0.3578] |
| | | Intron | 0.2963 [0.2762, 0.3159] | -0.1568 [-0.1757, -0.1380] | 0.1902 [0.1583, 0.2244] | -0.0337 [-0.0998, 0.0295] | -0.0031 [-0.0416, 0.0334] |
| | | Exon | -0.6837 [-0.7375, -0.6317] | 0.0110 [-0.0232, 0.0454] | -0.6211 [-0.7094, -0.5251] | -0.2393 [-0.4031, -0.0658] | -0.2604 [-0.3311, -0.1922] |
| | | Regulatory | -0.0435 [-0.0715, -0.0184] | 0.0352 [0.0051, 0.0679] | -0.1171 [-0.1616, -0.0741] | -0.3692 [-0.4592, -0.2897] | -0.0636 [-0.1237, 0.0096] |
| | Locus age * location | Intergenic | 0.0047 [0.0045, 0.0049] | 0.0059 [0.0057, 0.0061] | 0.0058 [0.0054, 0.0061] | 0.0098 [0.0088, 0.0108] | 0.0020 [0.0014, 0.0026] |
| | | Intron | 0.0016 [0.0014, 0.0018] | 0.0035 [0.0032, 0.0039] | 0.0027 [0.0023, 0.0031] | 0.0016 [0.0006, 0.0028] | 0.0014 [0.0006, 0.0021] |
| | | Exon | -0.0056 [-0.0060, -0.0052] | -0.0050 [-0.0053, -0.0048] | -0.0048 [-0.0055, -0.0038] | -0.0082 [-0.0108, -0.0056] | -0.0017 [-0.0023, -0.0010] |
| | | Regulatory | -0.0007 [-0.0009, -0.0004] | -0.0044 [-0.0047, -0.0040] | -0.0038 [-0.0041, -0.0033] | -0.0032 [-0.0044, -0.0022] | -0.0017 [-0.0026, -0.0010] |
| Impure | | mu | 19.3431 [19.0660, 19.5918] | 26.1808 [25.6007, 26.6928] | 24.4376 [24.0615, 24.8185] | 27.1998 [26.7889, 27.6029] | 28.0334 [27.6742, 28.4180] |
| | | Sample age | 0.0010 [0.0095, 0.0105] | 0.0141 [0.0136, 0.0145] | 0.0126 [0.0113, 0.0137] | 0.0113 [0.0097, 0.0128] | 0.0121 [0.0111, 0.0132] |
| | Location | Intergenic | 0.2920 [0.2350, 0.3454] | 0.0642 [0.0168, 0.1162] | 0.4437 [0.3491, 0.5292] | 1.3135 [1.1837, 1.4715] | 0.6328 [0.5605, 0.7099] |
| | | Intron | -0.0800 [-0.1363, -0.0248] | -0.2057 [-0.2759, -0.1493] | 0.0966 [0.0126, 0.1956] | 0.6052 [0.4593, 0.7528] | -0.1737 [-0.2633, -0.0934] |

| | | | | | | | |
|---|---|---|---|---|---|---|---|
| | | Exon | -0.2509<br>[-0.4003, -0.0894] | 1.7373<br>[1.6556, 1.8167] | 0.2298<br>[-0.0297, 0.4765] | -1.8676<br>[-2.3208, -1.4741] | -0.4591<br>[-0.6373, -0.2816] |
| | | Regulatory | 0.0388<br>[-0.0325, 0.1137] | -1.5957<br>[-1.6848, -1.5019] | -0.7701<br>[-0.8882, -0.6595] | -0.0511<br>[-0.2175, 0.1513] | -0.0000<br>[-0.1408, 0.1536] |
| | Locus age * location | Intergenic | 0.0080<br>[0.0075, 0.0086] | 0.0100<br>[0.0094, 0.0106] | 0.0109<br>[0.0096, 0.0121] | 0.0257<br>[0.0240, 0.0272] | 0.0203<br>[0.0190, 0.0216] |
| | | Intron | 0.0023<br>[0.0018, 0.0029] | 0.0047<br>[0.0037, 0.0056] | 0.0082<br>[0.0069, 0.0095] | 0.0044<br>[0.0024, 0.0062] | 0.0000<br>[-0.0014, 0.0018] |
| | | Exon | -0.0137<br>[-0.0152, -0.0124] | -0.0061<br>[-0.0067, -0.0056] | -0.0065<br>[-0.0101, -0.0032] | -0.0131<br>[-0.0173, -0.0088] | 0.0036<br>[0.0022, 0.0051] |
| | | Regulatory | 0.0033<br>[0.00265, 0.0039] | -0.0086<br>[-0.0096, -0.0076] | -0.0126<br>[-0.0141, -0.0113] | -0.0170<br>[-0.0191, -0.0148] | -0.0239<br>[-0.0266, -0.0215] |

Means and 95% highest posterior density intervals of parameters sampled from the posterior distribution of a generalized linear mixed model in which allele length is treated as dependent on sample, motif, surrounding sequence type (exon, intron, regulatory, or intergenic), estimated microsatellite locus age, and interaction between surrounding sequence type and locus age. Units for allele length and locus age are nucleotides and millions of years respectively. The effects of locus age and of the interaction term are therefore in nucleotides per million years.

**Supplementary Table 10: Collection and sequencing information for 26 modern Adélie penguin samples used in this study**

| Accession number | Collection date | Collection location | Latitude / Longitude | Sex | Material description | Collector (s) | Average sequencing depth |
|---|---|---|---|---|---|---|---|
| AP1.AP961174 | January 1997 | Torgerson Island, Antarctic Peninsula | 64° 46' S 64° 05' W | Undetermined | Seutin-preserved blood | Carol Vleck | 25.3 |
| AP2.AP961175 | January 1997 | Torgerson Island, Antarctic Peninsula | 64° 46' S 64° 05' W | Undetermined | Seutin-preserved blood | Carol Vleck | 27.2 |
| AP3.AP961178 | January 1997 | Torgerson Island, Antarctic Peninsula | 64° 46' S 64° 05' W | Undetermined | Seutin-preserved blood | Carol Vleck | 23.5 |
| AP4.AP961184 | January 1997 | Torgerson Island, Antarctic Peninsula | 64° 46' S 64° 05' W | M | Seutin-preserved blood | Carol Vleck | 25.7 |
| AP5.AP961194 | January 1997 | Torgerson Island, Antarctic Peninsula | 64° 46' S 64° 05' W | Undetermined | Seutin-preserved blood | Carol Vleck | 24.7 |

| | | | | | | | |
|---|---|---|---|---|---|---|---|
| AP6.AP961205 | January 1997 | Torgerson Island, Antarctic Peninsula | 64° 46’ S 64° 05’ W | Undetermined | Seutin-preserved blood | Carol Vleck | 20.1 |
| B2a | 2014 | Mawson region | 67° 36’ S 62° 52’ E | Undetermined | soft tissue in EtOH | Jane Younger, Louise Emmerson | 18.6 |
| B3a | 2014 | Mawson region | 67° 36’ S 62° 52’ E | Undetermined | soft tissue in EtOH | Jane Younger, Louise Emmerson | 20.8 |
| B4a | 2014 | Mawson region | 67° 36’ S 62° 52’ E | Undetermined | soft tissue in EtOH | Jane Younger, Louise Emmerson | 17.5 |
| B5a | 2014 | Mawson region | 67° 36’ S 62° 52’ E | Undetermined | soft tissue in EtOH | Jane Younger, Louise Emmerson | 18.2 |
| CA1.T401 | 2002/2003 | Cape Adare | 71° 17’ S 170° 14’ E | Undetermined | Seutin-preserved blood | David Lambert, John Macdonald | 25.8 |
| CA3.T415 | 2002/2003 | Cape Adare | 71° 17’ S 170° 14’ E | Undetermined | Seutin-preserved blood | David Lambert, John Macdonald | 19.9 |
| CA4.T403 | 2002/2003 | Cape Adare | 71° 17’ S 170° 14’ E | Undetermined | Seutin-preserved blood | David Lambert, John Macdonald | 19.1 |

| | | | | | | | |
|---|---|---|---|---|---|---|---|
| CA5.T409 | 2002/2003 | Cape Adare | 71° 17’ S<br>170° 14’ E | Undetermined | Seutin-preserved blood | David Lambert, John Macdonald | 23.0 |
| CB1.T526 | 2001 | Cape Bird | 77° 10’ S<br>166° 41’ E | F | Seutin-preserved blood | Craig Millar, Peter Ritchie | 21.7 |
| CB2.T528 | 2001 | Cape Bird | 77° 10’ S<br>166° 41’ E | F | Seutin-preserved blood | Craig Millar, Peter Ritchie | 23.9 |
| CB4.T525 | 2001 | Cape Bird | 77° 10’ S<br>166° 41’ E | F | Seutin-preserved blood | Craig Millar, Peter Ritchie | 19.9 |
| CB5.T527 | 2001 | Cape Bird | 77° 10’ S<br>166° 41’ E | M | Seutin-preserved blood | Craig Millar, Peter Ritchie | 17.9 |
| CLA02_11 | 2002/2003 | Coulman Island | 73° 29’ S<br>169° 45’ E | Undetermined | Seutin-preserved blood | David Lambert, John Macdonald | 19.1 |
| CLA02_13 | 2002/2003 | Coulman Island | 73° 29’ S<br>169° 45’ E | Undetermined | Seutin-preserved blood | David Lambert, John Macdonald | 22.5 |
| CLA_18 | 2002/2003 | Coulman Island | 73° 29’ S<br>169° 45’ E | Undetermined | Seutin-preserved blood | David Lambert, John Macdonald | 24.2 |

| | | | | | | | |
|---|---|---|---|---|---|---|---|
| CLA_20 | 2002/2003 | Coulman Island | 73° 29’ S<br>169° 45’ E | Undetermined | Seutin-preserved blood | David Lambert, John Macdonald | 22.5 |
| II1.T1 | 2002/2003 | Inexpressible Island | 74° 53’ S<br>163° 45’ E | Undetermined | Seutin-preserved blood | David Lambert, John Macdonald | 19.6 |
| II2.T3 | 2002/2003 | Inexpressible Island | 74° 53’ S<br>163° 45’ E | Undetermined | Seutin-preserved blood | David Lambert, John Macdonald | 22.5 |
| II4.T2 | 2002/2003 | Inexpressible Island | 74° 53’ S<br>163° 45’ E | Undetermined | Seutin-preserved blood | David Lambert, John Macdonald | 21.6 |
| II5.T15 | 2002/2003 | Inexpressible Island | 74° 53’ S<br>163° 45’ E | Undetermined | Seutin-preserved blood | David Lambert, John Macdonald | 22.6 |

**Supplementary Table 11: Collection information for 21 ancient Adélie samples used in this study**

| Accession | Collection Location | Latitude and Longitude Locality | Estimated Age (ybp) | Material Description | Collector | Average Sequencing Depth | Average read length (bp) |
|---|---|---|---|---|---|---|---|
| CB070106.05 | Inexpressible Island CBS18/06 | 74° 53’ S 163° 45’ E | 4670[1] | bone | Baroni / Salvatore | 5.2 | 48.6 |
| CB070108.07 | Inexpressible Island CBS 25/2006 | 74° 53’ S 163° 45’ E | 4140[1] | bone | Baroni / Salvatore | 8.1 | 43.5 |
| CB070117.03 | N Adélie Cove | 74° 46’ S 164° 0’ E | 6220[1] | bone | Baroni / Salvatore | 8.2 | 56.0 |
| CB070121.08 | Dunlop Island CBS 37/06 | 77° 14’ S 163° 30’ E | 30,000[2] | Bone | Baroni / Salvatore | 5.7 | 52.3 |
| CB070121.13 | Dunlop Island CBS 38/06 | 77° 14’ S 163° 30’ E | 25,600[2] | Bone | Baroni / Salvatore | 8.3 | 53.9 |
| CB070121.16 | Dunlop Island CBS 38/06 | 77° 14’ S 163° 30’ E | **46,587** | Bone | Baroni / Salvatore | 6.4 | 77.3 |

| | | | | | | | |
|---|---|---|---|---|---|---|---|
| CB111216.01 | Cape Bird - McDonald Beach CBS 05/2011 | 77° 15' S 166° 23' E | **3505** | bone and soft tissue | Baroni / Millar / Salvatore / Lorenzini | 11.2 | 65.7 |
| CB111216.02 | Cape Bird - McDonald Beach c/o CBS 05/2011 | 77° 15' S 166° 23' E | 3505[1] | bone | Baroni / Millar / Salvatore / Lorenzini | 10.5 | 69.9 |
| CB111216.05 | Cape Bird - McDonald Beach c/o CBS 05/2011 | 77° 15' S 166° 23' E | 3485[1] | bone | Baroni / Millar / Salvatore / Lorenzini | 7.8 | 67.3 |
| CB111216.06 | Cape Bird - McDonald Beach c/o CBS 05/2011 | 77° 15' S 166° 23' E | 3485[1] | bone | Baroni / Millar / Salvatore / Lorenzini | 7.7 | 68.8 |
| CB111216.07 | Cape Bird - McDonald Beach c/o CBS 05/2011 | 77° 15' S 166° 23' E | **3492** | bone | Baroni / Millar / Salvatore / Lorenzini | 10.6 | 62.7 |
| CB111217.10 | Cape Bird - McDonald Beach CBS 07/2011 | 77° 15' S 166° 23' E | 3434[1] | bone | Baroni / Millar / Salvatore / Lorenzini | 7.8 | 61.9 |
| CB111217.11 | Cape Bird - McDonald Beach CBS 07/2011 | 77° 15' S 166° 23' E | **3377** | bone | Baroni / Millar / Salvatore / Lorenzini | 6.1 | 51.5 |
| CB111228.08 | Cape Royds c/o CBS 12/2011 | 77° 33' S 166° 10' E | **2619** | bone | Baroni / Millar / Salvatore / Lorenzini | 6.4 | 59.5 |
| CB111229.15 | Cape Royds c/o CBS 15/2011 | 77° 33' S 166° 10' E | **716** | bone | Baroni / Millar / Salvatore / Lorenzini | 7.9 | 58.7 |

| | | | | | | | |
|---|---|---|---|---|---|---|---|
| CB111231.04 | Cape Barne - Sunk Lake CBS 18/2011 | 77° 15’ S 166° 23’ E | 329[1] | bone | Baroni / Millar / Salvatore / Lorenzini | 5.1 | 84.8 |
| CB121212.12 | Spike Cape CBS2/2012 | 77° 18’ S 163° 34’ E | **5388** | bone | Baroni / Millar / Salvatore | 7.1 | 41.3 |
| CB130104.09 | Marble Point (CBS 12/2012) | 77° 25’ S 163° 49’ E | 2886[1] | bone | Baroni / Millar / Salvatore / Parks | 7.2 | 68.8 |
| CB130105.05 | Marble Point (CBS 14/2012) | 77° 25’ S 163° 49’ E | **2886** | bone and soft tissue | Baroni / Millar / Salvatore / Parks | 10.2 | 55.9 |
| CB130105.13 | Marble Point (CBS 15/2012) | 77° 25’ S 163° 49’ E | 2886[1] | bone | Baroni / Millar / Salvatore / Parks | 7.7 | 60.3 |
| CB130106.05 | Marble Point (CBS 16/2012) | 77° 25’ S 163° 49’ E | **2556** | bone | Baroni / Millar / Salvatore / Parks | 11.5 | 56.2 |
| CB130106.24 | Marble Point (CBS 19/2012) | 77° 25’ S 163° 49’ E | 2249[1] | bone | Baroni / Millar / Salvatore / Parks | 7.4 | 55.7 |
| CB130106.26 | Marble Point (CBS 19/2012) | 77° 25’ S 163° 49’ E | **1937** | bone | Baroni / Millar / Salvatore / Parks | 7.5 | 53.5 |

Estimated ages in bold are from direct $^{14}C$ dating of samples, remaining ages estimated from $^{14}C$ dating of one or more samples collected at same locale and from similar stratigraphy[1], or stratigraphic dating of soil samples[2]. Dates of collection are represented in accession names as CB*YYMMDD*.##.

**Supplementary Table 12. Genomes used**

| Species | Genome size (Gb) | % aligned | % chicken aligned | % coding | Reference |
|---|---|---|---|---|---|
| **Zebra finch** | 1.256 | 68.25 | 73.10 | 1.91 | [1] |
| **Medium ground-finch** | 1.073 | 70.05 | 66.25 | 2.06 | [2] |
| **American crow** | 1.095 | 71.07 | 68.18 | 2.05 | [2] |
| **Golden-collared manakin** | 1.160 | 67.66 | 68.54 | 1.82 | [2] |
| **Rifleman** | 1.045 | 73.03 | 66.66 | 1.73 | [2] |
| **Budgerigar** | 1.133 | 70.03 | 69.24 | 1.89 | [3] |
| **Kea** | 1.062 | 76.19 | 70.09 | 1.72 | [2] |
| **Peregrine falcon** | 1.174 | 74.68 | 74.67 | 2.10 | [4] |
| **Red-legged seriema** | 1.146 | 76.03 | 73.86 | 1.54 | [2] |
| **Carmine bee-eater** | 1.067 | 74.55 | 68.77 | 1.54 | [2] |
| **Downy woodpecker** | 1.175 | 56.99 | 58.61 | 1.84 | [2] |
| **Rhinoceros hornbill** | 1.106 | 72.46 | 69.48 | 1.58 | [2] |
| **Bar-tailed trogon** | 1.086 | 73.77 | 69.65 | 1.56 | [2] |
| **Cuckoo-roller** | 1.148 | 75.37 | 73.87 | 1.59 | [2] |
| **Speckled mousebird** | 1.086 | 72.41 | 68.76 | 1.48 | [2] |
| **Barn owl** | 1.139 | 75.54 | 73.23 | 1.47 | [2] |
| **Bald eagle** | 1.280 | 69.56 | 75.02 | 1.75 | [2] |

| | | | | | |
|---|---|---|---|---|---|
| **White-tailed eagle** | 1.145 | 76.70 | 74.45 | 1.51 | 2 |
| **Turkey vulture** | 1.180 | 75.15 | 74.81 | 1.27 | 2 |
| **Dalmatian pelican** | 1.169 | 75.35 | 74.48 | 1.49 | 2 |
| **Little egret** | 1.212 | 71.86 | 77.25 | 1.74 | 2 |
| **Crested ibis** | 1.240 | 72.00 | 75.50 | 1.82 | 2 |
| **Great cormorant** | 1.155 | 74.80 | 73.58 | 1.46 | 2 |
| **Northern fulmar** | 1.153 | 76.55 | 74.78 | 1.52 | 2 |
| **Adélie penguin** | 1.251 | 71.09 | 75.06 | 1.69 | 2 |
| **Emperor penguin** | 1.283 | 69.84 | 75.44 | 1.74 | 2 |
| **Red-throated loon** | 1.145 | 76.58 | 74.35 | 1.46 | 2 |
| **White-tailed tropicbird** | 1.167 | 74.16 | 73.64 | 1.56 | 2 |
| **Sunbittern** | 1.099 | 75.05 | 71.10 | 1.51 | 2 |
| **Killdeer** | 1.225 | 71.70 | 74.51 | 1.81 | 2 |
| **Grey-crowned crane** | 1.139 | 76.22 | 73.92 | 1.58 | 2 |
| **Hoatzin** | 1.209 | 70.49 | 72.92 | 1.73 | 2 |
| **Annas hummingbird** | 1.115 | 67.91 | 66.25 | 1.98 | 2 |
| **Chimney swift** | 1.125 | 69.57 | 68.02 | 1.92 | 2 |
| **Chuck-wills widow** | 1.145 | 73.31 | 71.61 | 1.50 | 2 |
| **MacQueens bustard** | 1.095 | 76.62 | 71.66 | 1.55 | 2 |
| **Red-crested turaco** | 1.176 | 72.94 | 73.10 | 1.57 | 2 |
| **Common cuckoo** | 1.159 | 69.74 | 69.96 | 1.91 | 2 |
| **Brown mesite** | 1.101 | 74.33 | 70.74 | 1.62 | 2 |

| | | | | | |
|---|---|---|---|---|---|
| **Yellow-throated sandgrouse** | 1.098 | 75.74 | 71.29 | 1.55 | 2 |
| **Pigeon** | 1.112 | 73.97 | 70.98 | 2.03 | 5 |
| **American flamingo** | 1.145 | 76.64 | 74.21 | 1.44 | 2 |
| **Great crested grebe** | 1.151 | 74.14 | 72.71 | 1.37 | 2 |
| **Chicken** | 1.127 | | | 2.09 | 6 |
| **Turkey** | 1.080 | 79.29 | 76.52 | 1.93 | 7 |
| **Peking duck** | 1.105 | 78.03 | 73.13 | 2.57 | 8 |
| **White-throated tinamou** | 1.060 | 67.34 | 62.46 | 1.91 | 2 |
| **Common ostrich** | 1.228 | 69.74 | 72.07 | 1.69 | 2 |
| **Green anole lizard** | 1.835 | 44.26 | 10.41 | | 9 |
| **Human** | 3.200 | 2.96 | 8.41 | | 10 |
| **Chimpanzee** | 3.374 | 3.50 | 10.48 | | 11 |
| **Orangutan** | 3.516 | 3.35 | 10.47 | | 12 |
| **Mouse** | 2.780 | 3.09 | 7.66 | | 13 |
| **Rat** | 2.891 | 2.36 | 6.08 | | 14 |
| **Guinea pig** | 2.778 | 3.84 | 9.52 | | 15 |
| **Horse** | 2.534 | 2.75 | 6.19 | | 16 |
| **Opossum** | 3.678 | 2.69 | 8.72 | | 17 |
| **Platypus** | 2.037 | 5.29 | 9.53 | | 18 |
| **Western clawed frog** | 1.542 | 3.57 | 4.93 | | 19 |
| **Zebrafish** | 1.810 | 2.66 | 4.28 | | 20 |
| **Fugu** | 0.409 | 7.49 | 2.72 | | 21 |

| | | | | | |
|---|---|---|---|---|---|
| **Sea lamprey** | 1.048 | 1.90 | 1.75 | | [22] |
| **Lancelet** | 0.945 | 2.05 | 1.72 | | [23] |

Genome sizes are shown in Gb, along with the percentage of the genome present in the pairwise alignment with the chicken genome, the percentage of the chicken genome present in the same alignment, and the percentage of the genome accounted for by protein-coding sequence.

**Supplementary Table 13. TRF alignment score thresholds.**

| Period | Alignment score |
|---|---|
| 2 | 22 |
| 3 | 28 |
| 4 | 28 |
| 5 | 32 |
| 6 | 34 |

Taken from Willems *et al.* (2014) [24].

**Supplementary References**


1 Warren, W. C. *et al.* The genome of a songbird. *Nature* **464**, 757-762, doi:10.1038/nature08819 (2010).

2 Zhang, G. *et al.* Comparative genomics reveals insights into avian genome evolution and adaptation. *Science* **346**, 1311-1320, doi:10.1126/science.1251385 (2014).

3 Ganapathy, G. *et al.* High-coverage sequencing and annotated assemblies of the budgerigar genome. *GigaScience* **3**, 11, doi:10.1186/2047-217X-3-11 (2014).

4 Zhan, X. *et al.* Peregrine and saker falcon genome sequences provide insights into evolution of a predatory lifestyle. *Nat. Genet.* **45**, 563-566, doi:10.1038/ng.2588 (2013).

5 Shapiro, M. D. *et al.* Genomic diversity and evolution of the head crest in the rock pigeon. *Science* **339**, 1063-1067 (2013).

6 International Chicken Genome Sequencing Consortium. Sequence and comparative analysis of the chicken genome provide unique perspectives on vertebrate evolution. *Nature* **432**, 695-716 (2004).

7 Dalloul, R. A. *et al.* Multi-platform next-generation sequencing of the domestic Turkey (*Meleagris gallopavo*): Genome assembly and analysis. *PLoS Biol.* **8**, e1000475, doi:10.1371/journal.pbio.1000475 (2010).

8 Huang, Y. *et al.* The duck genome and transcriptome provide insight into an avian influenza virus reservoir species. *Nat. Genet.* **45**, 776-783, doi:10.1038/ng.2657 (2013).

9 Alföldi, J. *et al.* The genome of the green anole lizard and a comparative analysis with birds and mammals. *Nature* **477**, 587-591, doi:10.1038/nature10390 (2011).

10 International Human Genome Sequencing Consortium. Initial sequencing and analysis of the human genome. *Nature* **409**, 860-921 (2001).

11 The Chimpanzee Sequencing and Analysis Consortium. Initial sequence of the chimpanzee genome and comparison with the human genome. *Nature* **437**, 69-87, doi:10.1038/nature04072 (2005).

12 Locke, D. P. *et al.* Comparative and demographic analysis of orang-utan genomes. *Nature* **469**, 529-533, doi:10.1038/nature09687 (2011).

13 Mouse Genome Sequencing Consortium. Initial sequencing and comparative analysis of the mouse genome. *Nature* **420**, 520-562 (2002).

14 Rat Genome Sequencing Project Consortium. Genome sequence of the Brown Norway rat yields insights into mammalian evolution. *Nature* **428**, 493-521 (2004).

15 Lindblad-Toh, K. *et al.* A high-resolution map of human evolutionary constraint using 29 mammals. *Nature* **478**, 476-482, doi:10.1038/nature10530 (2011).

16 Wade, C. M. *et al.* Genome sequence, comparative analysis, and population genetics of the domestic horse. *Science* **326**, 865-867, doi:10.1126/science.1178158 (2009).

17 Mikkelsen, T. S. *et al.* Genome of the marsupial *Monodelphis domestica* reveals innovation in non-coding sequences. *Nature* **447**, 167-177, doi:10.1038/nature05805 (2007).

18 Warren, W. C. *et al.* Genome analysis of the platypus reveals unique signatures of evolution. *Nature* **453**, 175-183, doi:10.1038/nature07253 (2008).

19 Hellsten, U. *et al.* The genome of the Western clawed frog *Xenopus tropicalis*. *Science* **328**, 633-636, doi:10.1126/science.1183670 (2010).

20 Howe, K. *et al.* The zebrafish reference genome sequence and its relationship to the human genome. *Nature* **496**, 498-503, doi:10.1038/nature12111 (2013).

21 Kai, W. *et al.* Integration of the genetic map and genome assembly of fugu facilitates insights into distinct features of genome evolution in teleosts and mammals. *Genome Biol. Evol.* **3**, 424-442, doi:10.1093/gbe/evr041 (2011).

22 Smith, J. J. *et al.* Sequencing of the sea lamprey (*Petromyzon marinus*) genome provides insights into vertebrate evolution. *Nat. Genet.* **45**, 415-421, doi:10.1038/ng.2568 (2013).

23 Putnam, N. H. *et al.* The amphioxus genome and the evolution of the chordate karyotype. *Nature* **453**, 1064-1071, doi:10.1038/nature06967 (2008).

24 Willems, T. F., Gymrek, M., Highnam, G., Mittelman, D. & Erlich, Y. The landscape of human STR variation. *Genome Res.*, 1894-1904, doi:10.1101/gr.177774.114 (2014).
